# Effects of pathogen sexual reproduction on the evolutionary and epidemiological control provided by deployment strategies for two major resistance genes in agricultural landscapes

**DOI:** 10.1101/2023.02.02.526796

**Authors:** Marta Zaffaroni, Loup Rimbaud, Jean-François Rey, Julien Papaïx, Frédéric Fabre

## Abstract

- Resistant cultivars are of value for protecting crops from disease, but can be rapidly overcome by pathogens. Several strategies have been proposed to delay pathogen adaptation (evolutionary control), while maintaining effective protection (epidemiological control). Resistance genes can be *i*) combined in the same cultivar (pyramiding), *ii*) deployed in different cultivars sown in the same field (mixtures) or in different fields (mosaics), or *iii*) alternated over time (rotations). The outcomes of these strategies have been investigated principally in pathogens displaying pure clonal reproduction, but sexual reproduction may promote the emergence of superpathogens adapted to all the resistance genes deployed.
- We improved the spatially explicit stochastic model *landsepi* to include pathogen sexual reproduction, and then investigate the effect of sexual reproduction on evolutionary and epidemiological outcomes across deployment strategies for two major resistance genes.
- Sexual reproduction only favours the establishment of a superpathogen when single mutant pathogens are present together at a sufficiently high frequency, as in mosaic and mixture strategies.
- We concluded that, although sexual reproduction may promote the establishment of a superpathogen, it did not affect the optimal strategy recommendations for a wide range of mutation probabilities, associated fitness costs, and landscape organisations (notably the cropping ratio of resistant fields).

## 1 Introduction

The deployment of resistant cultivars in agricultural landscapes is a relatively low-input and cost-effective way to protect crops from plant pathogens. However, resistant cultivars have often been rapidly overcome by pathogens, especially when a single resistant cultivar is widely cultivated over a large geographic area (McDonald and Linde, 2002; Parlevliet, 2002; García-Arenal and McDonald, 2003). Ultimately, this may result in recurrent cycles of resistance deployment followed by rapid pathogen adaptation, often described as boom-and-bust cycles (McDonald and Linde, 2002). Several strategies have been proposed to promote a more durable management of resistant cultivars. These strategies involve increasing cultivated host genetic diversity (McDonald, 2010, 2014; Zhan et al., 2015) with the aim of confronting pathogens with eco-evolutionary challenges to prevent or delay their adaptation to plant resistance (evolutionary control), while maintaining effective disease protection (epidemiological control). Plant breeders can stack resistance sources in the same cultivar by pyramiding (McDonald and Linde, 2002; Fuchs, 2017), or farmers can alternate resistances over time by rotating cultivars in the same field (Curl, 1963). Host genetic diversity can also be introduced spatially. Resistant cultivars can be combined within the same field in cultivar mixtures (Wolfe, 1985; Mundt, 2002) or cultivated in different fields in landscape mosaics (Burdon et al., 2014; Zhan et al., 2015).

Given the multitude of deployment options, it is not straightforward to compare deployment strategies for identification of the optimal deployment strategy in a given epidemiological context. In addition, evolutionary and epidemiological control may not necessarily be correlated: any strategy designed to control the emergence of resistance-adapted pathogens in agro-ecosystems may potentially come into conflict with epidemiological control (Burdon et al., 2014; Papäix et al., 2018; Rimbaud et al., 2018a). Finally, particularly for airborne plant pathogens, which often disperse over large distances, deployment strategies are more likely to be effective if implemented across landscapes at large spatial scales, rendering experimental testing logistically demanding (but see Lohaus et al. 2000; Zhu et al. 2000; Djian-Caporalino et al. 2014; Koller et al. 2018). Many mathematical models have been developed to overcome these difficulties, to facilitate assessments of the variation of evolutionary and epidemiological outcomes across different resistance deployment strategies (reviewed by Rimbaud et al. 2021). These models have been used to unravel the effects of resistance deployment strategies on pathogen epidemiology and evolution, and to compare these strategies in a given epidemiological context.

Most of the models reviewed by Rimbaud et al. (2021) include only selection and/or mutation as evolutionary forces. This approach is suitable for the simulation of pathogens with purely clonal reproduction systems. Under the hypothesis of a purely clonal reproduction system, new pathogen variants are already present (possibly at low frequency) at the beginning of the simulated period, are introduced through migration, or are generated by mutation. However, some pathogens are not purely clonal and their life cycles include at least one sexual event per cropping season (mixed reproduction system), with some even reproducing exclusively by sexual means (purely sexual reproduction system). Of the 43 plant pathogens analysed by McDonald and Linde (2002), only 17 have exclusively clonal reproduction, the other 26 pathogens presenting at least one sexual reproduction event during their life cycle. The genetic recombination occurring during sexual reproduction can efficiently create gene combinations that would be accessible only through sequential mutation events in a purely clonal reproduction system. Several authors have argued that pathogens with mixed reproduction system have the highest potential for evolving and breaking down the resistances deployed in agriculture (McDonald and Linde, 2002; Stam and McDonald, 2018). Genetic recombination first creates many new variants of the pathogen (Tibayrenc and Ayala, 2002; Halkett et al., 2005). The populations of the fittest variants then expand rapidly through clonal reproduction, potentially breaking down the resistance, (*i*.*e*. increasing the frequency of pathogen strains adapted to the resistance genes present). Genetic recombination can, therefore, have a major impact on the evolutionary and epidemiological outcomes of resistance deployment strategies (Arenas et al., 2018; Stam and McDonald, 2018). It has been shown that even low rates of recombination in pests and pathogens have profound implications for policies concerning drug and pesticide resistance (Halkett et al., 2005). Similarly, by mixing the genotypes of parental individuals, recombination can favour the emergence of the generalist superpathogens able to overcome pyramided cultivars (McDonald and Linde, 2002; Uecker, 2017). However, the ability of recombination to favour the emergence of superpathogens also depends on subtle interactions between mutation and recombination rates on the one hand, and pathogen population size on the other (Althaus and Bonhoeffer, 2005). Indeed, recombination can generate variants accumulating infectivities, but it can also break down such such genetic combinations (Hadany and Beker, 2003).

Despite the potentially major impact of the pathogen reproduction system on the epidemiological and evolutionary control provided by resistance deployment strategies, this impact has been little studied and is poorly understood. Genetic recombination is considered in only three (Sapoukhina et al., 2009; Xu, 2012; Crété et al., 2020) of the 69 models reviewed by Rimbaud et al. (2021) and in a recent study by Saubin et al. (2021). These studies considered pathogens with mixed reproduction systems, but they did not compare purely clonal reproduction with mixed reproduction systems, all other things being equal. It is, therefore, difficult to assess the impact of reproduction system on the epidemiological and evolutionary control provided by resistance deployment strategies from the data currently available. In addition, these works focused on just one or two resistance deployment strategies, preventing a global assessment of all possible spatiotemporal deployment options. They highlighted the role of the fitness cost of resistance in superpathogen persistence (Xu, 2012), and in the efficacy of rotation (Crété et al., 2020) and mixture (Xu, 2012; Sapoukhina et al., 2009) strategies. In addition, Saubin et al. (2021) assessed the impact of ploidy on resistance durability, revealing that resistance durability was greater, but more variable, for diploid pathogens.

Here, we investigated the effect of pathogen sexual reproduction on the evolutionary and epidemiological control achieved with four main categories of deployment strategies (rotation, pyramiding, mixture and mosaic). We adapted the *landsepi* model (Rimbaud et al., 2018b), which simulates the spread of epidemics across an agricultural landscape and the evolution of a pathogen in response to the deployment of host resistance, to include pathogen sexual reproduction. We then used this model to compare the resistance deployment strategies considered for situations in which two major resistance genes conferring immunity are deployed. The new model is flexible enough to vary resistance deployment strategy and pathogen life cycle, making it possible to compare pathogens with different reproduction systems (purely clonal *vs*. mixed). We parameterised the model to simulate grapevine downy mildew, which is caused by the oomycete *Plasmopara viticola*. However, our general conclusions are likely to have broader implications to other pathosystems.

## 2 Description

### 2.1 Model overview

The model used in this study is an adapted version of that presented by Rimbaud et al. (2018b), which simulates the clonal reproduction, spread and evolution of a pathogen in an agricultural landscape over multiple cropping seasons. Here, we introduce between-season sexual reproduction to address the issue of pathogens with mixed reproduction systems. Multiple clonal reproduction events occur during the life cycle of these pathogens, with a final sexual reproduction event at the end of the host cropping season. We split the modelled cropping season into two different time periods: *i*) within the cropping season, when multiple clonal reproduction events take place, and *ii*) the period between cropping seasons, when a single sexual reproduction event may take place. Below, we describe only the major changes between cropping seasons, the modifications within cropping seasons being only minor. The entire model is described in *Supporting Information* note S1, and the code is available from the R package *landsepi* (v1.2.4, Rimbaud et al. 2022).

### 2.2 Landscape and resistance deployment strategies

We considered agricultural landscapes randomly generated with a T-tessellation algorithm (Papäix et al., 2014) in which four cultivars were randomly allocated to fields: a susceptible cultivar (SC) initially infected with a pathogen not adapted to any resistance, two resistant cultivars, each carrying a single resistance gene (RC_1_ and RC_2_), and one resistant cultivar carrying both resistance genes (RC_12_). We first allocated a proportion 1 *− ϕ*_1_ of fields to receive SC, the remaining *ϕ*_1_ candidate fields then being allocated a cultivar according to one of the following strategies:

1. Mosaics: RC_1_ and RC_2_ are cultivated in the equal proportions of the candidate fields (*ϕ*_2_ = 0.5);
2. Mixture: both RC_1_ and RC_2_ are cultivated in all the candidate fields, in equal proportions within each field (*ϕ*_2_ = 0.5);
3. Rotations: RC_1_ and RC_2_ are cultivated alternately in candidate fields for a fixed number of cropping seasons (three-year rotation).
4. Pyramiding: RC_12_ is cultivated in all candidate fields.

A cultivar carrying a major resistance gene is assumed to be immune to disease (*i*.*e*. pathogen infection rate is equal to 0), unless the pathogen has acquired the corresponding infectivity gene, according to the so-called “gene-for-gene” hypothesis (Leonard, 1977; Thompson and Burdon, 1992). A non-adapted pathogen (denoted “WT” here for “wild type”) can acquire infectivity gene *g ∈ {*1, 2*}* through a single mutation, with a probability *τ*_*g*_, or, alternatively, through sexual reproduction with another individual pathogen carrying such an infectivity gene. Infectivity genes confer an ability to break down the associated major gene resistance on the pathogen. The evolution of infectivity may be penalised by a fitness cost (*θ*_*g*_) on susceptible hosts (Brown, 2015; Laine and Barr’es, 2013; Thrall and Burdon, 2003). This fitness cost is represented in the model as a lower infection rate for mutant pathogens on hosts not carrying the corresponding resistance gene. Here, a pathogen genotype is represented by a set of binary variables indicating whether it carries infectivity genes able to overcome cultivar resistance genes. There are four possible pathogen genotypes: wild-type, unable to break down the resistance conferred by any resistance gene (“00”), single mutant “SM_1_” (or “SM_2_”), able to break down to the first (or second) resistance gene (“10” and “01”, respectively), and superpathogen “SP”, able to break down both resistance genes (“11”). The relative infection rates of these pathogens on the different cultivars are summarised in Table 1.

**Table 1:** Plant-pathogen interaction matrix.

| | | Host genotype $v$ | | | |
| --- | --- | --- | --- | --- | --- |
|  |  | SC | RC<sub>1</sub> | RC<sub>2</sub> | RC<sub>12</sub> |
| Pathogen genotypes $p$ | WT | 1 | 0 | 0 | 0 |
| | SM<sub>1</sub> | $1-\theta_1$ | 1 | 0 | 0 |
| | SM<sub>2</sub> | $1-\theta_2$ | 0 | 1 | 0 |
| | SP | $(1-\theta_1)(1-\theta_2)$ | $1-\theta_2$ | $1-\theta_1$ | 1 |

The matrix gives the coefficient by which the infection rate is multiplied. The value of this coefficient reflects the relative infection rates for the wild-type (WT) and adapted (single mutants SM<sub>1</sub> and SM<sub>2</sub>, and SP) pathogen genotypes on the susceptible (SC) and resistant cultivars carrying a single major resistance gene (cultivar RC<sub>1</sub> and cultivar RC<sub>2</sub>), or their combination (RC<sub>12</sub>). $\theta_1$ and $\theta_2$ are the fitness costs of adaptation with respect to the major resistance genes considered.

### 2.3 Demogenetic dynamics within the cropping season

The demogenetic dynamics of the host-pathogen interaction within the cropping season are based on a compartmental model with a discrete time step, schematically reported in Fig. 1. Below, H_i,v,t_, L_i,v,p,t_, I_i,v,p,t_, R_i,v,p,t_, and P_i,p,t_ denote the numbers of healthy, latent, infectious and removed individuals, and of pathogen propagules, respectively, in the field *i* = 1,…,J, for cultivar *v* = 1,…, V, pathogen genotype *p* = 1,…,P at time step t=1,…,T*×*Y (Y is the number of cropping seasons and T the number of time steps per season). Note that, in this model, an “individual” is defined as a given amount of plant tissue, and is referred to as a “host” hereafter for the sake of simplicity. At the beginning of the cropping season, healthy hosts are contaminated with the primary inoculum generated at the end of the previous cropping season.

**Figure 1:**
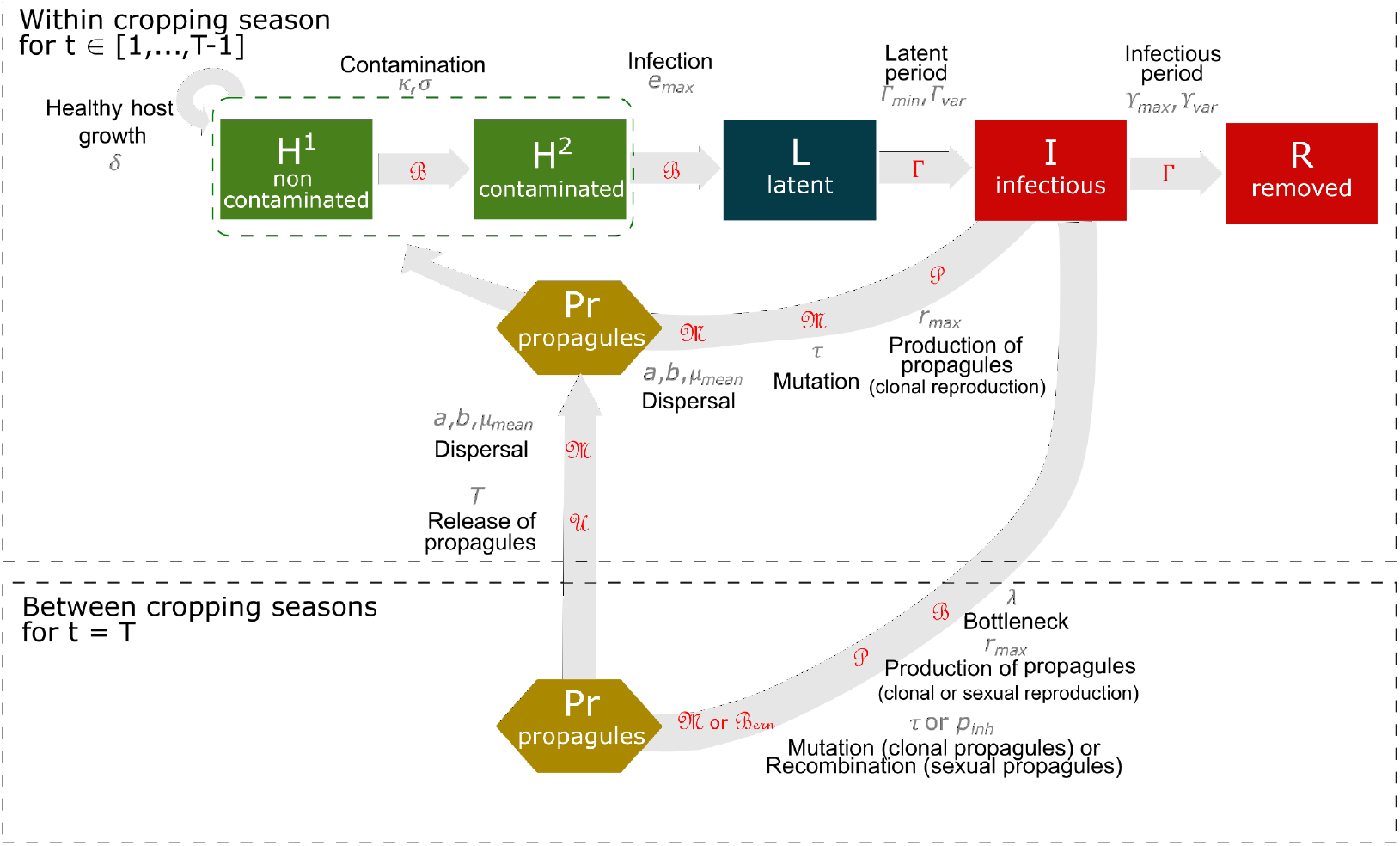
Model overview. Within-cropping season dynamics: healthy hosts can be contaminated by pathogen propagules (produced both at the end of the previous cropping season and within the current cropping season) and may become infected. Following a latent period, infectious hosts start producing propagules through clonal reproduction. These propagules may mutate and disperse across the landscape. At the end of the infectious period, infected hosts become epidemiologically inactive. Qualitative resistance prevents the infection of contaminated hosts, *i*.*e*. their transition to the latently infected state. Green boxes indicate healthy hosts contributing to host growth, as opposed to diseased plants (*i*.*e*. symptomatic, red boxes) or plants with latent infections (dark blue box). Between-cropping season dynamics: at the end of each cropping season, pathogens experience a bottleneck during the off-season period, and propagules are then produced (by clonal or sexual reproduction). Clonal propagules may mutate, whereas genetic recombination may occur during sexual reproduction. Propagules produced between host cropping seasons are gradually released during the following host cropping season. The parameters associated with epidemiological processes are indicated in grey and detailed in Table 2. The distributions used to simulate stochasticity in model transitions are indicated in red; ℬ: binomial, Γ: gamma, 𝒫: Poisson, ℳ: multinomial, 𝒰: uniform, ℬ*ern*: Bernoulli. Host logistic growth is deterministic. The entire model is described in *Supporting Information* note S1.

### 2.4 Demogenetic dynamics between cropping seasons

The demogenetic dynamics of the host-pathogen interaction between cropping seasons is presented schematically in Fig. 1. At the end of the cropping season, the crop is harvested and the leaves of the host plants fall to the ground, imposing a potential bottleneck on the pathogen population before the start of the next cropping season. The remaining hosts produce clonal or sexual propagules. Clonal propagules can mutate in the same way as they do during the cropping season. The production of propagules through sexual reproduction and the possibility of genetic recombination are detailed in the section 2.4.1. The propagules produced during the period between cropping seasons, whether clonal or sexual, are uniformly released throughout the following cropping season, constituting the primary inoculum.

**Table 2:** Summary of model parameters and numerical simulation plan (factors in bold are varied according to a complete factorial design).

| Notation | Parameter | Value | Source |
| --- | --- | --- | --- |
| Simulation Factors |  |  |  |
| Y | Number of cropping seasons | 50 years | Fixed |
| T | Number of time steps in a cropping season | 120 days | Fixed |
| **J** | **Number of fields in the landscape** | **[155; 154; 152; 153; 156]** | **Varied** |
| V | Number of host cultivars | [2<sup>a</sup>, 3<sup>b</sup>] | Fixed |
| Initial conditions and seasonality (same value for all cultivars) |  |  |  |
| $C_v^0$ | Plantation host density of cultivar v (in pure crops) | 1 m<sup>-2</sup> | Fixed |
| $C_v^{max}$ | Maximal host density of cultivar v (in pure crops) | 20 m<sup>-2</sup> | Fixed |
| $\delta_v$ | Host growth rate of cultivar v | 0.1 day<sup>-1</sup> | [1] |
| $\Phi$ | Initial probability of infection of susceptible hosts | 5.10<sup>-4</sup> | Fixed |
| $\lambda$ | Off-season survival probability of pathogen spores | 10<sup>-4</sup> | Fixed |
| Pathogen aggressiveness components |  |  |  |
| $e_{max}$ | Maximal expected infection rate | 0.9 spore<sup>-1</sup> | [2,3] |
| $\Gamma_{min}$ | Minimal expected latent period duration | 7 days | See note S2 |
| $\Gamma_{var}$ | Variance of the latent period duration | 8 days | See note S2 |
| $\Upsilon_{max}$ | Maximal expected infectious period duration | 14 days | See note S2 |
| $\Upsilon_{var}$ | Variance of infectious period duration | 22 days | See note S2 |
| $r_{max}$ | Maximal expected propagule production rate | 2 spores.day<sup>-1</sup> | See note S2 |
| Sexual reproduction |  |  |  |
| **R** | **Pathogen reproduction system** | **[purely clonal, mixed]** | **Varied** |
| $p_{inh}$ | Probability of a sexual propagule inheriting the genotype at locus $g$ from parent $Par_1$ genotype | 0.5 | Fixed |
| Pathogen dispersal |  |  |  |
| $g(\cdot)$ | Dispersal kernel | Power-law function | See note S1 |
| $\mu_{mean}$ | Mean dispersal distance | 20 m | [4] |
| $a$ | Scale parameter | 40 | [4] |
| $b$ | Width of the tail | 7 | [4] |
| Contamination of healthy hosts |  |  |  |
| $\pi(\cdot)$ | Contamination function | Sigmoid | See note S1 |
| $\sigma$ | Related to the position of the inflection point | 3 | [4] |
| $\kappa$ | Related to the position of the inflection point | 5.33 | [4] |
| Host-pathogen genetic interaction |  |  |  |
| G | Total number of major genes | 2 | Fixed |
| $\tau_g$ | **Mutation probability for infectivity gene g** | **[10<sup>-7</sup>; 10<sup>-4</sup>]** | **Varied** |
| $\rho_g$ | Efficiency of major gene g | 1 | |
| $\theta_g$ | **Cost of infectivity of infectivity gene g** | **[0; 0.25; 0.5]** | **Varied** |
| Landscape organisation |  |  |  |
|  | **Resistance deployment strategy** | **MIxture; MOsaic; PYramiding; ROtation** | **Varied** |
| $\alpha$ | Level of spatial aggregation | 0 | Fixed |
| $\varphi_1$ | **Cropping ratio of fields in which resistance is deployed** | **[0.17; 0.33; 0.5; 0.67; 0.83]** | **Varied** |
| $\varphi_2$ | Relative cropping ratio of RC<sub>2</sub> | 0.5<sup>c</sup> | Fixed |

<sup>a</sup> : pyramiding; <sup>b</sup> : mixture, mosaic, rotation; <sup>c</sup> : for mixture and mosaic only. Source: [1] Bove and Rossi (2020); [2] Bove et al. (2019); [3] Boso and Kassemeyer (2008); [4] Rimbaud et al. (2018b).

#### 2.4.1 Pathogen sexual reproduction

In field *i*, the pool of infectious hosts associated with the same cultivar *v* undergoes sexual reproduction. Two parental infectious hosts, infected with pathogens *Par*_1_ and *Par*_2_, respectively, are randomly sampled without replacement from the pool of infectious hosts. The *c* = *{Par*_1_; *Par*_2_*}* pair produces 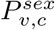 propagules, drawn from a Poisson distribution in which the expectation is the sum of the number *r*_*max*_ of propagules produced by each of the parental infectious hosts:

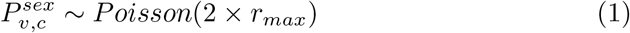

The genotype of each propagule is then retrieved from the parental genotypes: the genotype at every locus *g* is randomly sampled from one of the two parents *{Par*_1_; *Par*_2_*}*. For example, assuming that parental infection *Par*_1_ provides infectivity genes against resistance gene *g* = 1 (corresponding to genotype “10”) and parental infection *Par*_2_ provides infectivity genes effective against resistance *g* = 2 (genotype “01”), the resulting propagule genotype may be the same as that of one of the two parents (with probability 0.5), an SP genotype “11” (probability 0.25), or a WT genotype “00” (probability 0.25). This process is iterated for all the pairs *c* = 1,…, *C* of infectious hosts associated with all the cultivars *v* = 1,…, *V* in a given field *i*, resulting in a total number of sexual propagules:

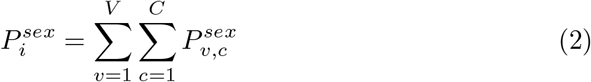

### 2.5 Propagule dispersal

Clonal and sexual propagules disperse similarly (no dispersal dimorphism) within the landscape according to a power-law dispersal kernel.

### 2.6 Simulation plan and model outputs

#### 2.6.1 Model parameterisation for *Plasmopara viticola*

We parameterised the model to simulate epidemics of *Plasmopara viticola*, the causal agent of grapevine downy mildew, which has a mixed reproduction system (Wong et al., 2001; Gessler et al., 2011). Downy mildew is a real threat to grapevines in all vine-growing areas of the world, causing significant yield losses and leading to the massive use of pesticides (Gessler et al., 2011). In recent years, breeders have been developing programs for breeding resistance to grapevine downy mildew, resulting in the creation of several resistant varieties, with the aim of lowering rates of fungicide application on grapevines. However, *P. viticola* has already been shown to have a high evolutionary potential, as demonstrated by the rapid emergence of fungicide resistance (Blum et al., 2010; Chen et al., 2007) and the breakdown of some of the resistances deployed (Peressotti et al., 2010; Delmas et al., 2016; Paineau et al., 2022). All the model parameters used in the simulations are listed in Table 2.

#### 2.6.2 Simulation plan

The model is used to assess evolutionary and epidemiological outputs for different deployment strategies. In addition to the four resistance deployment strategies considered (mosaic, mixture, rotation, pyramiding), we varied the cropping ratio of fields where resistance is deployed (*ϕ*_1_, five values), while assuming similar relative proportions of the two resistant cultivars (*ϕ*_2_ = 0.5 in mixtures and mosaics). We simulated different pathogen evolutionary potentials, by varying the mutation probability (*τ*, two levels) and the fitness cost (*θ*, three values) while assuming the same characteristics for both major genes (*i*.*e. τ*_*g*_ = *τ* and *θ*_*g*_ = *θ ∀g ∈* 1, 2). We explored the effect of the pathogen reproduction system by either having the pathogen reproduce sexually at the end of the cropping season (mixed reproduction system) or having no sexual reproduction event (purely clonal reproduction system). The abovementioned factors were explored with a complete factorial design of 240 parameter combinations (Table 2). Simulations were also performed with five different landscape structures (with about 155 fields and a total area of 2 *×* 2 km^2^, see Fig. S11 in the *Supporting Information*) and 48 replications in each landscape structure, resulting in a total of 240 replicates per parameter combination. The whole numerical design represents a total of 57600 simulations. Each simulation was run for 50 cropping seasons of 120 days each. Trial simulations showed that this simulation horizon was sufficiently long to differentiate between deployment strategies in terms of their evolutionary and epidemiological performances.

#### 2.6.3 Model outputs

At the end of a simulation run, the results were evaluated by considering evolutionary and epidemiological outputs. For evolutionary outputs, we determined the time point at which the generalist superpathogen SP was established in the resistant host population. We first studied SP establishment by defining *E*_*SP*_ a binary variable set to 1 if the SP becomes established before the end of a simulation run and 0 otherwise. Assuming that the SP became established, we then studied the time to establishment *T*_*SP*_ . This time corresponds to the time point at which the number of resistant host plants infected with SP exceeds a threshold above which extinction in a constant environment becomes unlikely. We also determined the time required for the two single mutants to become established (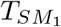 and 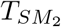). Finally, we monitored the size of the superpathogen population 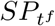 and the maximum number of heterogeneous parental pairs 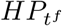 (*i*.*e*. parental pairs involving *SM*_1_ and *SM*_2_) in the landscape after the bottleneck. In a given field and for a given host cultivar, the maximum number of heterogeneous parental pairs was calculated as the minimum between the population size of *SM*_1_ and *SM*_2_ after harvest at *t*^*f*^ ; which gives, for the whole landscape: 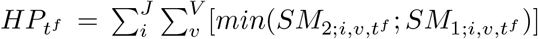. For epidemiological output, we assessed the area under the disease progress curve (AUDPC) to measure disease severity over the whole landscape, averaged across all the simulated cropping seasons. AUDPC is normalised by dividing by mean disease severity in a fully susceptible landscape; its value therefore ranges from 0 (*i*.*e*. no disease) to 1 (*i*.*e*. disease severity identical to that in a fully susceptible landscape).

### 2.7 Statistical analysis

We first used a classification tree to determine how the factors of interest and their interactions affected the binary evolutionary output *E*_*SP*_ . We considered the following six factors as qualitative explanatory variables: resistance deployment strategy, cropping ratio, mutation probability and fitness cost of the infectivity genes, the pathogen reproduction system and landscape structure. We then fitted a logistic regression to assess the relationship between *E*_*SP*_ and the time elapsed between the establishment of the two single mutants 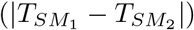, for a selected subset of factors. In addition, for each combination of resistance deployment strategy, mutation probability, fitness cost and pathogen reproduction system, we fitted second-order polynomial regressions (or second-order logistic regressions) to assess the response of *T*_*SP*_ and *AUDPC* (or *E*_*SP*_) to variations of cropping ratio. Note that fitting a second-order logistic regression was impossible for factor combinations that almost always or never led to SP establishment in the 240 replicates. In such cases, a secondorder polynomial regression was fitted instead. Finally, for each combination of resistance deployment strategy, mutation probability, fitness cost, pathogen reproduction system and cropping ratio, we fitted local polynomial regressions to the temporal dynamics of the population of 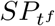 and 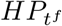 .

Statistical analyses were performed with R (v4.0.5, R Core Team 2021) software. The function *rpart* within the package *rpart* (v4.1.16, Therneau et al. 2022) was used to fit the classification and regression trees (we set a minimum number of values in any terminal node equal to 3% the total number of values). The function *glm* within the package *stats* (v3.6.2, R Core Team 2022) was used to fit the logistic regression (*glm*(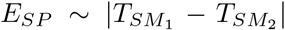 strategy, family = “binomial”). The function *geom smooth* within the package *ggplot2* (v3.3.6, Wickham et al. 2022) was used to fit second-order logistic (method = “*glm*”, formula = *y ∼* poly(*x*, 2), family = “binomial”), second-order polynomial (method = “lm”, formula = *y ∼* poly(*x*, 2)) and local polynomial (method = “loess”, formula = *y ∼ x*) regressions.

## 3 Results

The SP became established before the end of the 50-year simulation in 75.2 % of the 57600 simulations. In these 43320 simulations, the mean time to SP establishment was 4.69 years, and the 2.5th and 97.5th percentiles were 0.6 and 31.5 years, respectively. For the 57600 simulations performed, the AUDPC ranged from 14% (*i*.*e*., mild epidemics) to 99% (*i*.*e*., severe epidemics). Below, we determine the roles of the principal factors driving such variability in output.

### 3.1 Factors affecting superpathogen establishment

We constructed a classification tree for identifying parameter combinations leading to SP establishment (*E*_*SP*_) (Fig. 2A). *E*_*SP*_ was dependent principally on the mutation probability, the resistance deployment strategy and the fitness cost.

**Figure 2:**
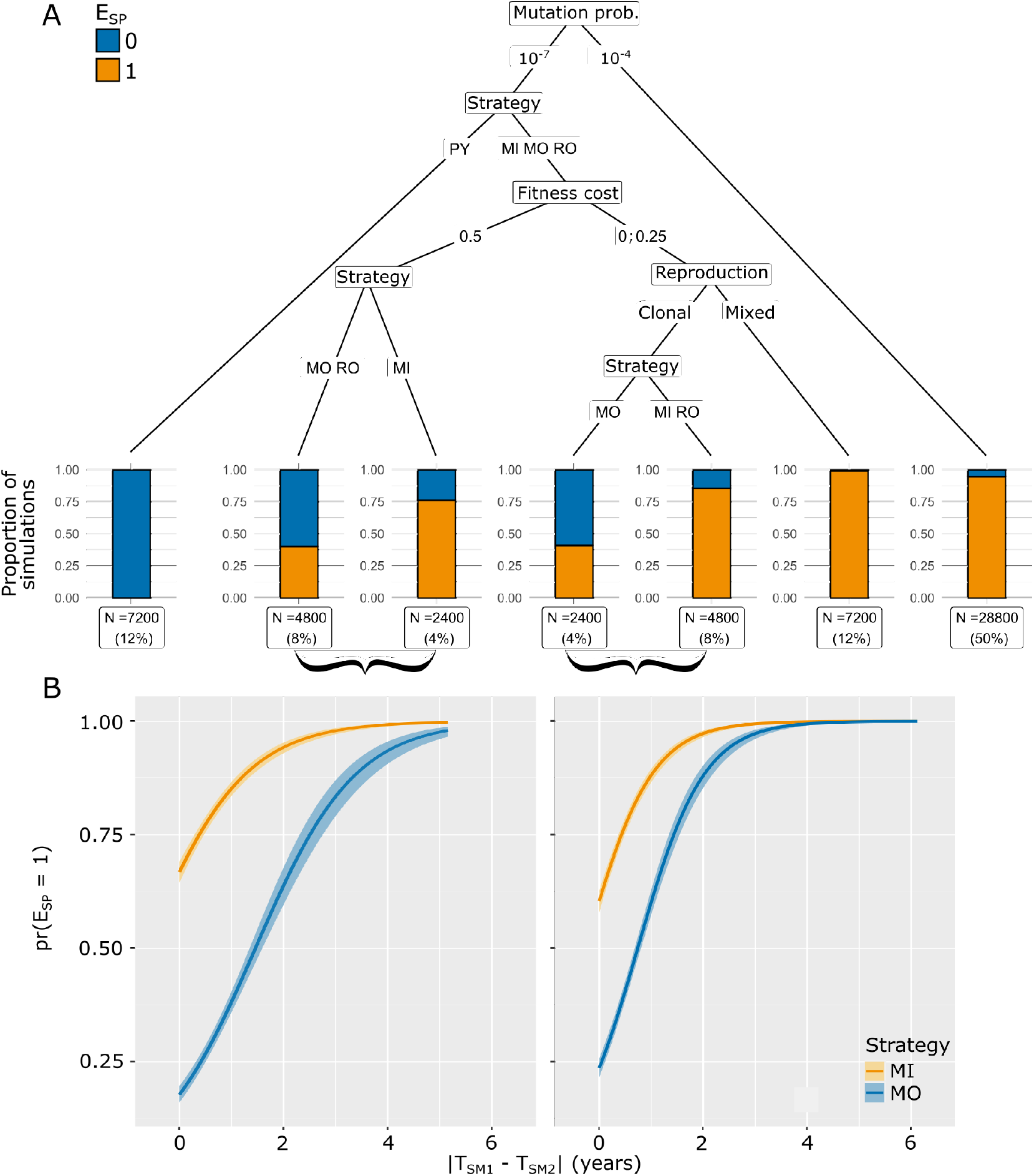
(A) Classification tree for the binary output *E*_*SP*_ . The number and proportion of simulations (of the 57600 performed) associated with each end node are indicated. Orange bars indicate the proportion of simulations in which the SP became established before the end of the simulation, whereas blue bars indicate the proportion of simulations in which this was not the case. The factors identified by the tree are the mutation probability for infectivity genes, the resistance deployment strategy (MIxture, MOsaic, ROtation and PYramiding), the fitness cost of infectivity genes and the pathogen reproduction system (purely clonal or mixed). (B) Relationship between the time elapsed between the establishment of the two single mutants (SM_1_ and SM_2_) and the probability of superpathogen emergence (*pr*(*E*_*SP*_ = 1)) for the MIxture and MOsaic strategies. Logistic regression was used to fit relationships to simulation outputs corresponding to the combination of parameters highlighted in brackets under the final nodes of the tree. Confidence intervals are delimited by the 2.5th and 97.5th percentiles.

At high mutation probabilities, the SP almost invariably became established in the pathogen population, regardless of the other factors. At low mutation probabilities, specific combinations of these factors determined whether or not the SP became established. For example, the SP was never established in conditions in which the resistance genes were pyramided in the same cultivar. The SP became established in less than one in two simulations when resistance genes were deployed in *i*) mosaic and rotation, for high fitness costs (*θ* = 0.5); *ii*) mosaic, for fitness costs below 0.5 and purely clonal reproduction. For the remaining parameter combinations, the SP became established in more than one in two simulations. The pathogen reproduction system had a secondary influence on SP establishment. However, for mixture, mosaic and rotation strategies with a low or no fitness cost, the SP almost always became established for pathogens with a mixed reproduction system, whereas the proportion of simulations in which the SP became established was substantially lower for pathogens with a clonal reproduction system, particularly for mosaic strategies.

At low mutation probabilities, SP establishment was a highly stochastic event in mixture, mosaic and rotation strategies; it occurred in 41% to 87% of the simulations, depending on the values of the other factors (Fig. 2A). We improved the resolution of the corresponding final nodes, by hypothesising, for mosaic and mixture strategies, that SP establishment was dependent on the time interval between the establishment of the two single mutants |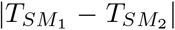 |. This hypothesis was based on the rationale that longer intervals would result in one of the two resistant hosts remaining an empty ecological niche for longer. It can, therefore, be infected by the SP if it emerges through mutation or recombination. This hypothesis holds only for the mosaic and mixture strategies, as the two resistant hosts must be deployed at the same time, excluding *de facto* the rotation strategies from the subsequent analysis. As expected, the probability of SP establishment increased sharply with 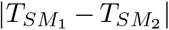, whatever the fitness cost. Moreover, the probability of SP establishment was systematically higher for mixtures than for mosaics (Fig. 2B). Finally, a specific feature of rotation strategies may also favour the emergence of the SP regardless of the pathogen reproduction system. Indeed, a SP generated by mutation from a single mutant late in the season (*i*.*e*. when the ecological niche is no longer empty) could still have an opportunity to establish itself in an empty niche if this event occurs shortly before the switch to a different variety in the rotation.

To deepen the analysis on the parameter combinations leading to SP establishment, we asses the relationship between the variable *E*_*SP*_ and the cropping ratio for all combinations of resistance deployment strategy, fitness cost and pathogen reproduction system considered (Fig. 3). We focused on low mutation probabilities, as shown in Fig. 3 (but see Fig. S1 for its analogous version with high mutation probability). The probability of *E*_*SP*_ generally increases with cropping ratio for mixture, mosaic and rotation strategies unless establishment is already certain at the lowest cropping ratio. However, for mixture strategies with non-zero fitness costs, the probability of *E*_*SP*_ for pathogens undergoing purely clonal reproduction follows a U-shaped curve, with the lowest probability of *E*_*SP*_ achieved for an intermediate cropping ratio. The SP was never established in simulations based on pyramiding strategies. Furthermore, for mixture and mosaic strategies, the probability of *E*_*SP*_ was consistently lower for pathogens with clonal rather than mixed reproduction. In addition, the probability of *E*_*SP*_ was lower for mosaics than for mixtures in pathogens with a clonal reproduction system.

**Figure 3:**
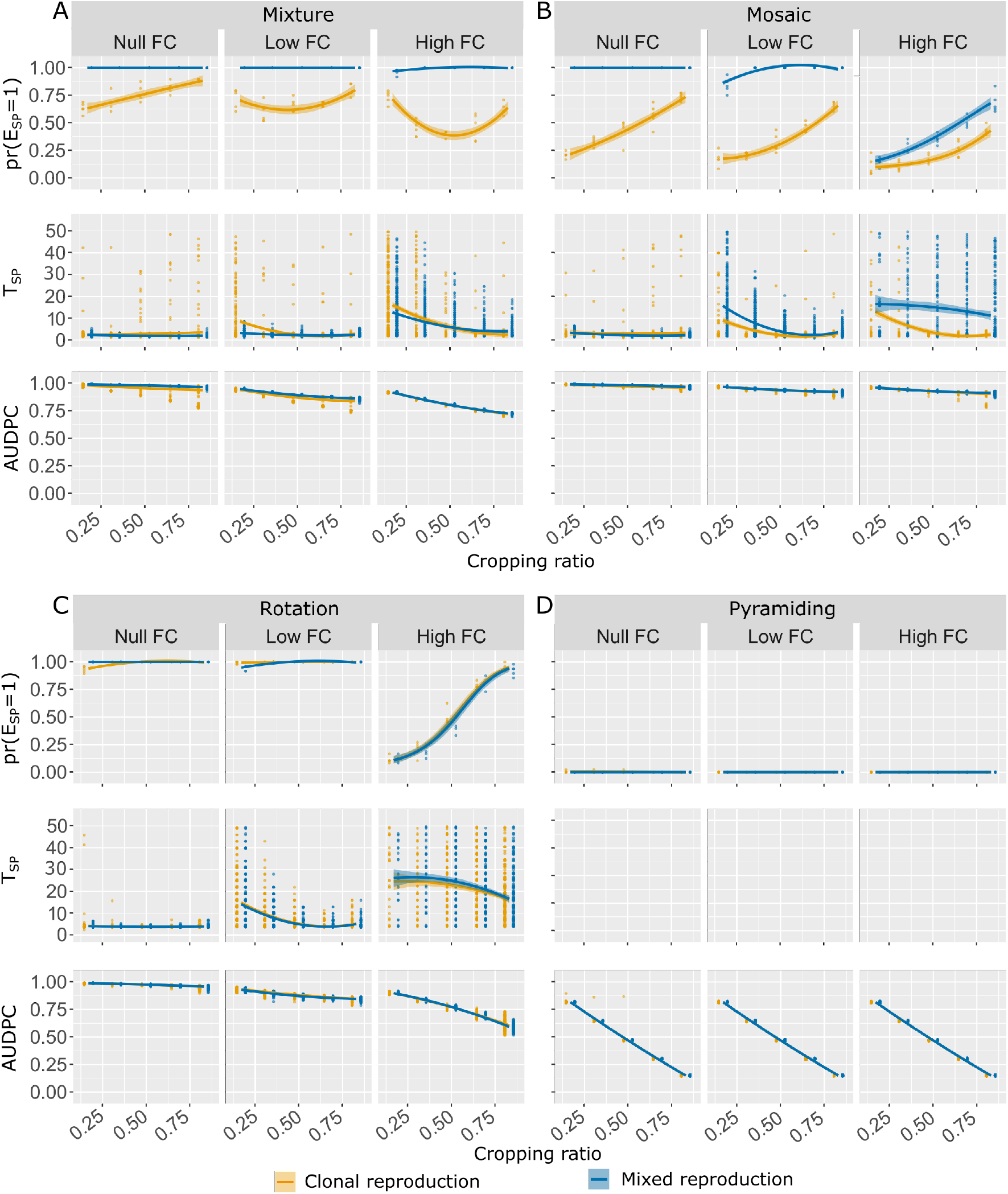
Probability of SP establishment (first row of each panel), time to SP establishment, given that the SP becomes established, (second row) and AUDPC (third row) at low (*τ* = 10^*−*7^) mutation probability and at zero (*θ* = 0), low (*θ* = 0.25) and high (*θ* = 0.5) fitness cost (FC). Panels show the effect on the probability of *E*_*SP*_, *T*_*SP*_, and AUDPC as a function of the cropping ratio for the two pathogen reproduction systems and the four deployment strategies considered. Curves are based on the fitting of logistic or second-order polynomial regressions to simulation outputs (represented by points, note that, in the first row of each panel, the points represent the proportion of *E*_*SP*_ = 1 among the 48 replicates); shaded envelopes delimited by the 2.5th and 97.5th percentiles.

The effect of the pathogen reproduction system on the probability of *E*_*SP*_ can be explained by the demogenetic dynamics of the pathogen population after the bottleneck at the end of the cropping season. Contrasting dynamics were, indeed, observed across resistance deployment strategies and fitness costs, as illustrated in Fig. 4 for intermediate cropping ratios. With mixture and mosaic strategies, the maximum number of heterogeneous parental pairs after the bottleneck 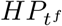 is relatively high, at least during the first 10 cropping seasons. In this setting, sexual recombination between single mutants favours the generation of SP propagules, which constitute the primary inoculum for the following season. Accordingly, the number of 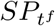 increases more rapidly, reaching a higher level for pathogens with mixed reproduction systems than for those with purely clonal reproduction, particularly if there is no fitness cost (for both mosaic and mixture strategies) or if the fitness cost is low (mixture strategy only). As a mirror effect, the number of 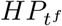 stabilises at lower levels for pathogens with a mixed reproduction system. This effect disappears at higher fitness costs. By contrast, the small number or absence of 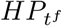 observed with the pyramiding and rotation strategies greatly decreases the likelihood of recombination between single mutants. Consequently, the production of SP propagules is not favoured by sexual reproduction in these strategies. Note that the trends in the demogenetic dynamics of 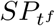 and 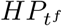 were similar for the other combinations of cropping ratios and mutation probabilities (Fig. S2-S10 in the *Supporting Information*).

**Figure 4:**
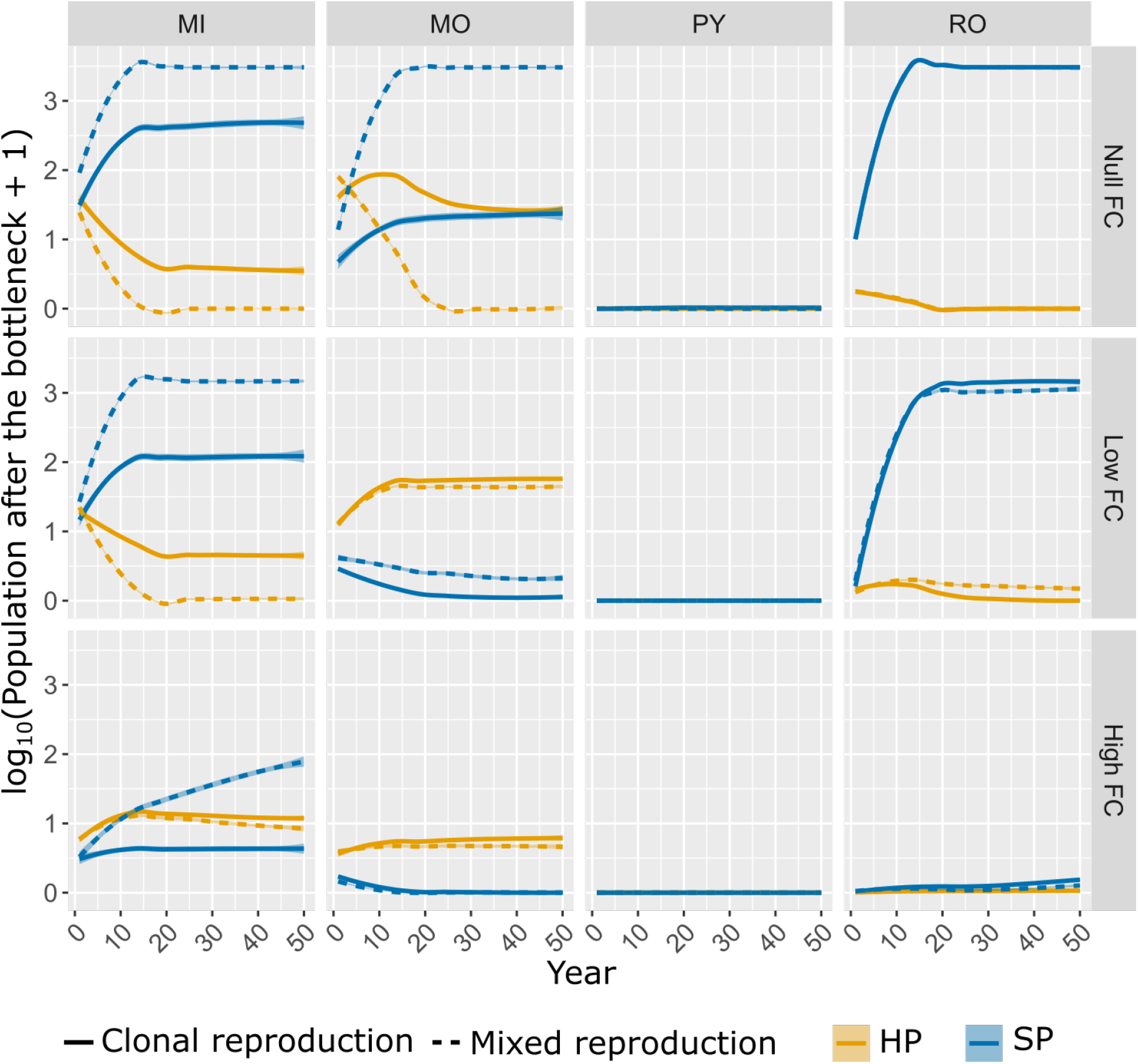
Population size of the superpathogen 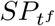 (in blue) and maximum number of heterogeneous parental pairs 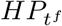 (in orange) in the landscape after the annual bottleneck. The curves represent the population dynamics across resistance deployment strategies (MIxture, MOsaic, ROtation and PYramiding), fitness costs and reproduction systems, at low mutation probability (*τ* = 10^*−*7^) and intermediate cropping ratio (*ϕ*_1_ = 0.5). The curves are based on the fitting of local polynomial regressions and shaded envelopes delimited by the 2.5th and 97.5th percentiles. Note that, at high fitness costs, the curves for pyramiding and rotation overlap.

### 3.2 Factors affecting the time to superpathogen establishment

The mean time to SP establishment *T*_*SP*_, estimated conditionally on SP establishment (*i*.*e*. for the subset of replicates such that *E*_*SP*_ = 1), generally decreases with cropping ratio. Furthermore, the type of reproduction does not generally influence *T*_*SP*_, except in the mosaic strategy (Fig. 3B). For this strategy, *T*_*SP*_ is lower for pathogens with purely clonal reproduction systems and non-zero fitness costs. However, at high mutation probability, *T*_*SP*_ is lower for pathogens with mixed rather than purely clonal reproduction systems, for fitness costs that are low or zero (Fig.S1 in the *Supporting Information*). Finally, our results show that the variance of *T*_*SP*_ increases substantially with fitness cost, suggesting that, in these contexts, the mean time to SP establishment poorly reflects the underlying evolutionary dynamics.

### 3.3 Factors affecting the mean area under the disease progress curve

In a fully susceptible landscape, the mean area under the disease progress curve, *AUDPC* _0_ was 0.63 for both pathogen reproduction systems. This value implies that diseased hosts (those in an infectious or removed state, see Fig. 1) accounted for a mean of 63% of the available host individuals over the entire period simulated. AUDPC generally decreased with cropping ratio (Fig. 3). At low mutation probability, the best epidemiological control (*i*.*e*. the lowest AU-DPC) was obtained with the pyramiding strategy, which decreased AUDPC by up to 86% at high cropping ratios, independently of the fitness cost incurred for pathogen adaptation. With the other strategies, the highest AUDPC reductions achieved (for the 240 replicates) were 22% for mosaics, 30% for mixtures, 49% for rotation. These values were obtained at a high cropping ratio and fitness cost. By contrast, almost no epidemic control (*i*.*e*. AUDPC *≈* 1) was observed for these strategies in the absence of a fitness cost. Finally, the pathogen reproduction system did not affect the AUDPC.

## 4 Discussion

We address the question of the effect of the type of pathogen reproduction system on the epidemiological and evolutionary control provided by plant resistance. Epidemiological control relates to plant health and the demographic dynamics of the pathogen, whereas evolutionary control relates to the durability of resistance and the genetic dynamics of the pathogen. Sexual reproduction principally favours the exchange of genes via recombination. We therefore studied the fate of the superpathogen during the deployment of two resistance genes.

### 4.1 Effect of the pathogen reproduction system on evolutionary and epidemiological outputs

McDonald and Linde (2002) hypothesised that pathogens with mixed reproduction systems pose the greatest risk of genetic resistance breakdown, because they benefit from the advantages of both reproduction systems. Betweencropping seasons, the occurrence of a single sexual reproduction event generates new pathogen genotypes that may combine mutations already present in the population. During the cropping season, clonal reproduction enable the fittest pathogen genotypes to invade the population rapidly. However, in tests of their risk model on 34 pathosystems, McDonald and Linde (2002) found no significant effects of the pathogen reproduction system on the risk of breakdown, which was instead affected by gene/genotype flow and mutation. Our results confirm the importance of mutation rate as a driver of pathogen evolution. Indeed, the SP was established in all simulations with a high mutation probability, regardless of the deployment strategy or pathogen reproduction system. This finding can be explained by the interplay between mutation probability and population size (Christiansen et al., 1998; Althaus and Bonhoeffer, 2005). Mean population size in this study was 1.3 *×* 10^7^. It follows that, at high mutation probability (*τ* = 10^*−*4^), at least one SP is likely to emerge through mutation during the first cropping season (which includes 17 clonal generations) in 89 of 100 simulations. Our results also show the effect of sexual reproduction on the likelihood of the generalist SP becoming established depends on the resistance deployment strategy. This finding goes a step further than the analysis presented by McDonald and Linde (2002), who did not consider the effect of deployment strategies. Our simulations suggest that recombination favours the establishment of the SP only when heterogeneous pairs of single mutant parents are potentially abundant after crop harvest. This is the case for the mosaic and mixture strategies (Fig. 4). For these strategies, populations of single mutant pathogens can increase in size on their specific hosts, with recombination subsequently occurring on susceptible hosts during sexual reproduction, potentially generating SP propagules between two cropping seasons. The timing of sexual reproduction is also a key element explaining why SP establishment is favoured by a mixed reproduction system. Indeed, the SP propagules generated by recombination during the offseason emerge right at the start of the following cropping season, when most hosts are healthy, favouring SP establishment in this empty ecological niche. By contrast, for pathogens with purely clonal reproduction, the SP is generated by mutation from a single mutant when the population is large enough. This event probably occurs late during the cropping season when the competition between the SP and the two single mutants for the infection of healthy hosts is much stronger. Accordingly, we found that the probability of SP establishment increased when the competition with the single mutants is lower, in particular when only one single mutant pathogen is established on a resistant host and the second host is free from disease (Fig. 2B).

By contrast, sexual reproduction does not favour the establishment of the SP in pyramiding and rotation strategies, because heterogeneous pairs of single mutants are scarce in these conditions (Fig. 4), as the cultivars carrying the single resistance genes are not deployed at all, or not deployed simultaneously. Similar result were reported in the context of the resistance to xenobiotics (Althaus and Bonhoeffer, 2005; Taylor and Cunniffe, 2022). In particular, sexual reproduction in fungi increases the frequency of the double-resistant strain adapted to a mixture of fungicides (as for the SP here) only when the frequency of single-resistant strains is significantly higher than that of double-resistant or avirulent strain (Taylor and Cunniffe, 2022).

### 4.2 No deployment strategy is universally optimal

Consistent with the findings of previous comparisons of deployment strategies (Djidjou-Demasse et al., 2017; Lof and van der Werf, 2017; Sapoukhina et al., 2009; Rimbaud et al., 2018a), our results confirm that no one strategy is universally optimal. Instead, the strategy used should be adapted to the pathosystem and production situation, and a decision must be taken as to whether to prioritise epidemiological or evolutionary outputs. With this in mind, given that pre-adapted pathogens were assumed to be initially absent, the order of magnitude of the mutation probability relative to pathogen population size is a key factor. Conversely, the pathogen reproduction system had no effect on strategy recommendations for various fitness costs, mutation probabilities and cropping ratios. Similarly Taylor and Cunniffe (2022) showed that sexual reproduction did not affect recommendations for the management of fungicides mixtures. At low mutation probabilities, a SP will emerge by mutation from the wildtype 1 in every 10000 times during the 17 *×* 50 generations within a simulation run. Providing that no preadapted pathogens are initially present, it explains the better performance of pyramiding over all other strategies (Leach et al., 2001). Pyramiding strategies ensure both epidemiological and evolutionary control of the targeted disease, as reported by Djian-Caporalino et al. (2014); Rimbaud et al. (2018a). In particular, the decrease in disease severity is proportional to the cropping ratio of the pyramided variety in the landscape as the dilution effect is maximal in this setting (Keesing and Ostfeld, 2021). For the other strategies, the probability of SP establishment generally increases with cropping ratio, as higher cropping ratios favour the development of large populations of single mutants, in turn favouring the emergence of the SP. However, for mixture strategies with fitness costs and pathogens with purely clonal reproduction, the relationship between cropping ratio and the probability of SP establishment is U-shaped. Among the mechanisms underlying this relationship, the intensity of the spill-over (*i*.*e*. infection of a new host from a reservoir population, Daszak et al. 2000) of simple mutants from fields cultivated with susceptible cultivar to fields cultivated with resistant cultivars should be a major driver. Indeed, the spill-over is maximum at intermediate cropping ratio. In this case, the population of simple mutants emerging from susceptible cultivars is more likely to quickly infect both resistant cultivars in the mixture, leaving few hosts for the SP, mutated from the SM, to infect. In the opposite, at either low or high cropping ratio, the spill-over of simple mutants from susceptible to resistant cultivar is reduced because adjacent fields are mostly sharing the same cultivar. It opens more rooms to the SP population to emerge from one of the two resistant cultivars in the mixture and to invade the other one.

At high mutation probabilities, the SP becomes established a mean of 1.2 years after the beginning of a simulation run for pyramiding strategies (Fig. S1D in the *Supporting Information*). There is no dilution effect at work during most of the 50-year time frame considered, and epidemiological and evolutionary control disappear. In this setting, the strategies delaying SP establishment for the longest were mosaic and rotation, at low cropping ratio and high fitness costs (Fig. S1B-C in the *Supporting Information*). With these strategies, the time to SP establishment decreased monotonically with cropping ratio. Higher fitness costs in these strategies also slowed SP establishment through disruptive selection. This mechanism exploits fitness costs to favour local host specialisation of the pathogen, limiting the likelihood of a generalist SP emerging (Barrett et al., 2009). Despite generally providing the best evolutionary control, the mosaic strategy was the worst strategy (in comparisons with rotation and mixture) in our conditions for epidemiological control. One key reason for this is the high probability of autoinfections, 0.82 on average, a consequence of our choice of large field sizes (mean of 160 m *×*160 m) relative to short mean pathogen dispersal distances (20 m). The frequent infection events resulting from propagules produced in the same field favours the mixture strategy over the mosaic strategy (Mundt, 2002). Like us, Djidjou-Demasse et al. (2017) also found that pyramiding and mosaic strategies provided similar levels of epidemiological control if the probability of autoinfection was high. In their study, frequent between-field infections and high rates of mutation were required for mosaic strategies to outperform pyramiding. Crucially, our results highlight the need for knowledge about mutation probability and the cost of infectivity to guide the choice of deployment strategy.

Unfortunately, there has been little quantitative characterization of these parameters (Laine and Barr’es, 2013). Point mutations are the simplest evolutionary events conferring virulence to a resistance gene. Such events occur once every 10^5^ to 10^7^ propagules per generation (Stam and McDonald, 2018). However, many other mutational events *sensu lato* (*e*.*g*. complete or partial gene deletion, insertion of transposable elements) increase the overall mutation probability conferring virulence (Daverdin et al., 2012). Unlike knowledge about the mutation probability, which can guide the choice as to whether or not to use a pyramiding strategy, the cost of infectivity has a monotonic influence: the higher the cost, the higher the levels of evolutionary and epidemiological control achieved. Such costs are not pervasive among plant-pathogenic fungi and vary with host genotype and abiotic environment (Laine and Barr’es, 2013). For example, substantial sporulation costs have been reported in rusts (Bahri et al., 2009; Thrall and Burdon, 2003) but no such costs evidenced for grapevine downy mildew (Toffolatti et al., 2012; Delmas et al., 2016)).

### 4.3 Further perspectives

The ecoevolutionary framework presented here represents a solid foundation for further investigations of the effects of other mechanisms linked to the sexual reproduction of pathogens. For example, we assume that all the sexual propagules emerge in the cropping season immediately following their production, but specialised reproductive structures can survive in the soil for many years (up to 5 years for *P. viticola*, Caffi et al. 2010). This feature may impact the outputs of deployment strategies, in particular rotations (Papavizas and Ayers, 1974). We also assume that sexual and clonal propagules have similar dispersal capacities. This may not always be the case, as shown for black sigatoka (Rieux et al., 2014) and grapevine downy mildew (Rossi and Caffi, 2012). Such dispersal dimorphism probably affects the effectiveness of resistance deployment strategies such as mixtures and mosaics (Papäix et al., 2018; Sapoukhina et al., 2010; Watkinson-Powell et al., 2020).

Furthermore, we focus here exclusively on qualitative resistance genes (*i*.*e*. major genes), but quantitative resistance is attracting increasing interest for use in pathogen control (Parlevliet, 2002; Niks et al., 2015). As the model can also handle quantitative resistances, it would be interesting to broaden our analysis in this direction. Recombination in a diverse pathogen population, as favoured by the partial effect of quantitative resistance on pathogens, might accelerate pathogen evolution towards higher levels of aggressiveness (Frézal et al., 2018; Drenth et al., 2019). Conversely, recombination, by breaking up blocks of coadapted genes, may slow the adaptation of pathogens to quantitative resistance genes (McDonald and Linde, 2002).

## Acknowledgements

This work was funded by the MEDEE project of the Ecophyto II APR Leviers Territoriaux (No.SIREPA 4621) national action plan.

## Data availability

Data sharing not applicable to this article as no datasets were generated or analysed during the current study. The code of the model is implemented in the R package *landsepi* : Landscape Epidemiology and Evolution (version 1.2.4, https://cran.r-project.org/web/packages/landsepi/index.html).

## Author contributions

M.Z, L.R, J.P., F.F. planned and designed the research. M.Z wrote the model. M.Z. and J.F.R. updated the *landsepi* package. M.Z. conducted the numerical experiment. M.Z, L.R, J.P., F.F. analyzed the numerical experiments and wrote the manuscript.

## Supporting Information

**Fig. S1** Probability of SP establishment, time before SP establishment and AUDPC at high mutation probability.

**Fig. S2-S10** Population size of the superpathogen and maximum number of heterogeneous parental pairs in the landscape after the bottleneck for combinations of mutation probabilities and cropping ratios.

**Fig. S11** The five landscapes considered in the simulation plan.

**Fig. S12** Distribution of the latent period duration of downy mildew caused by *Plasmopora viticola*.

**Fig. S13** Distribution of the infectious period duration of downy mildew caused by *Plasmopora viticola*.

**Table S1** Available data on the duration of latent and sporulation periods for downy mildew caused by *P. viticola* (and *formae speciales*).

**Note S1** Model equations.

**Note S2** parameterisation for *Plasmopara viticola*.

**Note S3** Calculation of the threshold for pathogen establishment considering sexual reproduction.

