## Supplementary Information for "Effects of pathogen sexual reproduction on the evolutionary and epidemiological control provided by deployment strategies for two major resistance genes in agricultural landscapes"

### Supporting Information

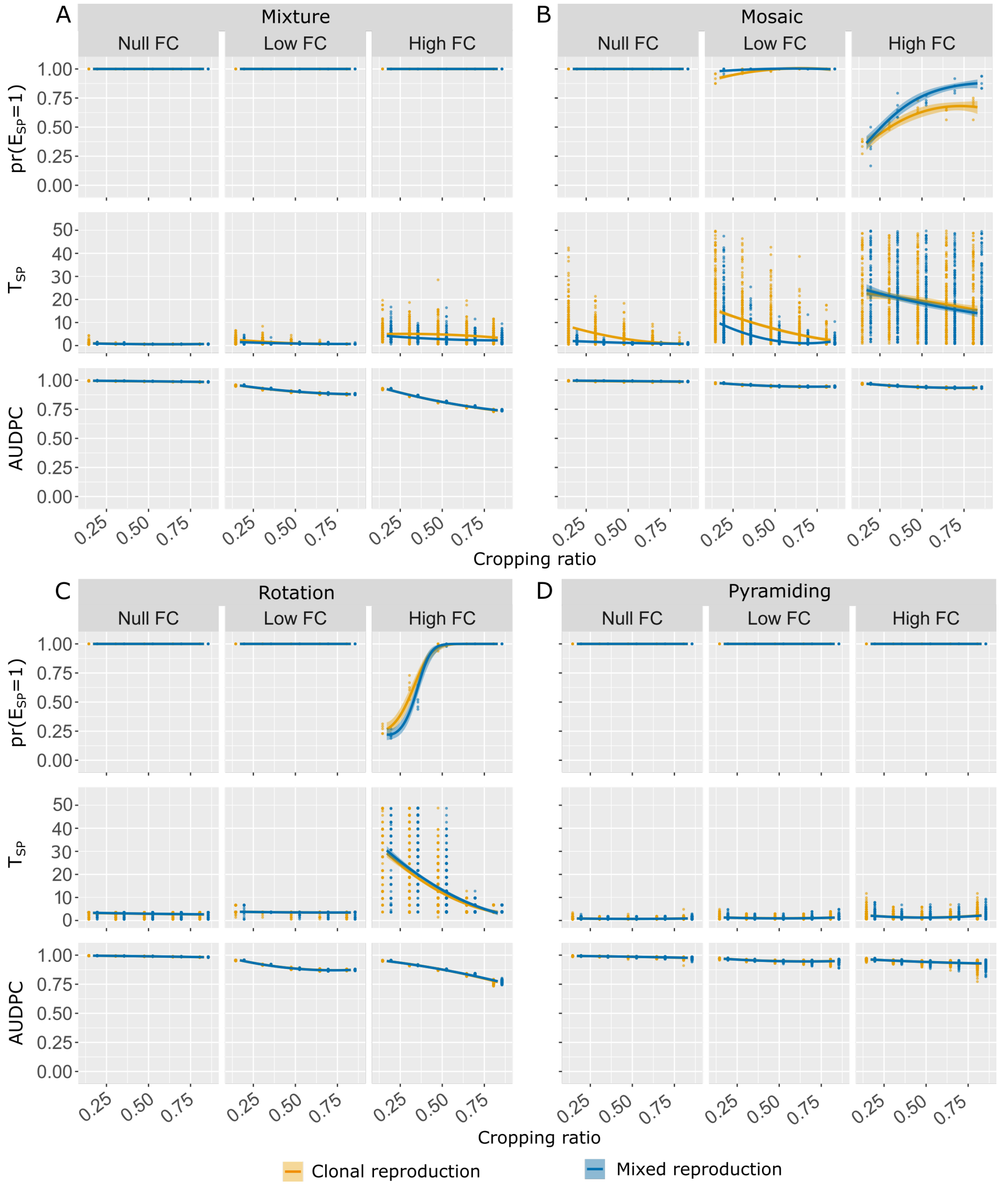


Figure S1: Probability of SP establishment (first row of each panel), time before SP establishment, given the SP gets established, (second row) and AUDPC(third row) at high ($\tau = 10^{-4}$) mutation probability and at null ($\theta = 0$), low ($\theta = 0.25$) and high ($\theta = 0.5$) fitness cost (FC). Panels show the effect on the probability of $E_{SP}$, $T_{SP}$, and AUDPC as a function of the cropping ratio for the two pathogen reproduction systems and the four deployment strategies considered. Curved are based on logistic or second order polynomial regression fitting performed on simulations outputs (represented by points, note that in the first row of each panel the points represent the proportion of $E_{SP} = 1$ among the 48 replicas), shaded envelopes are delimited by the 2.5th and 97.5th percentiles.

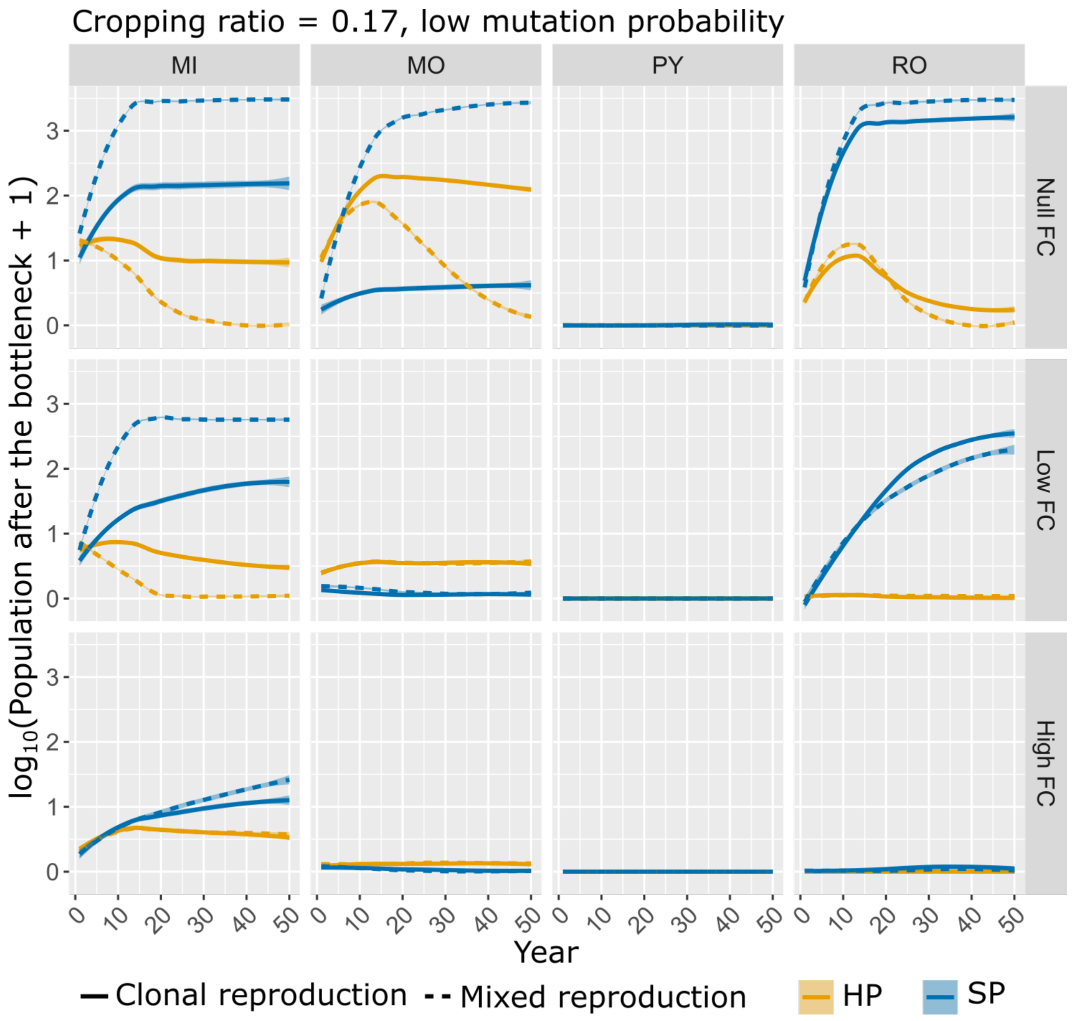


Figure S2: Population size of the superpathogen $SP_{tf}$ (in blue) and maximum number of heterogeneous parental pairs $HP_{tf}$ (in orange) in the landscape after the bottleneck. Curves represent populations dynamics across resistance deployment strategies (MIxture, MOsaic, ROtation and PYramiding) fitness costs and reproduction systems, at low mutation probability ($\tau = 10^{-7}$) and cropping ratio $\varphi_1 = 0.17$. Curves are based on the local polynomial regression fitting performed on simulations outputs. Shaded envelopes delimit the 2.5th and 97.5th percentiles.

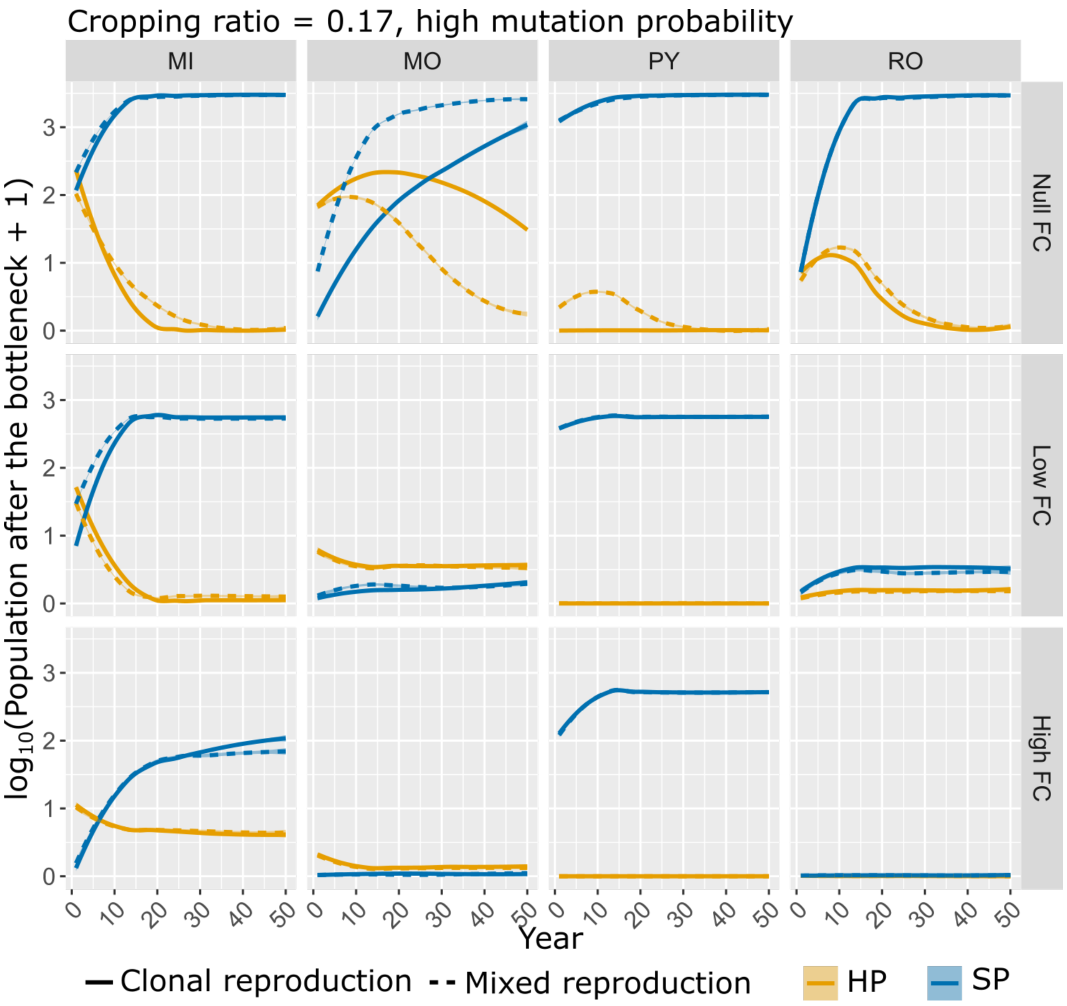


Figure S3: Population size of the superpathogen $SP_{t^f}$ (in blue) and maximum number of heterogeneous parental pairs $HP_{t^f}$ (in orange) in the landscape after the bottleneck. Curves represent populations dynamics across resistance deployment strategies (MIxture, MOsaic, ROtation and PYramiding) fitness costs and reproduction systems, at high mutation probability ($\tau = 10^{-4}$) and cropping ratio $\varphi_1 = 0.17$. Curves are based on the local polynomial regression fitting performed on simulations outputs. Shaded envelopes delimit the 2.5th and 97.5th percentiles.

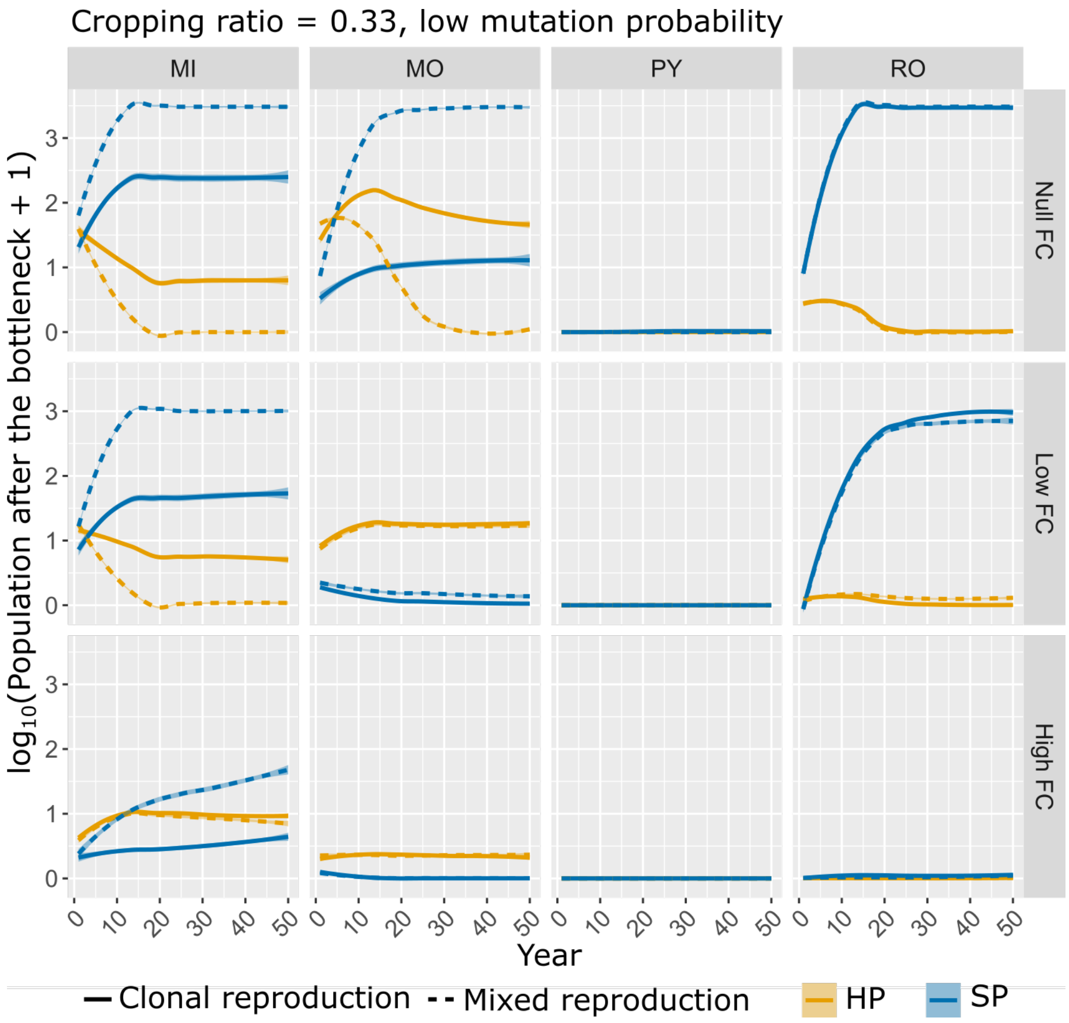


Figure S4: Population size of the superpathogen $SP_{tf}$ (in blue) and maximum number of heterogeneous parental pairs $HP_{tf}$ (in orange) in the landscape after the bottleneck. Curves represent populations dynamics across resistance deployment strategies (MIxture, MOsaic, ROtation and PYramiding) fitness costs and reproduction systems, at low mutation probability ($\tau = 10^{-7}$) and cropping ratio $\varphi_1 = 0.33$. Curves are based on the local polynomial regression fitting performed on simulations outputs. Shaded envelopes delimit the 2.5th and 97.5th percentiles.

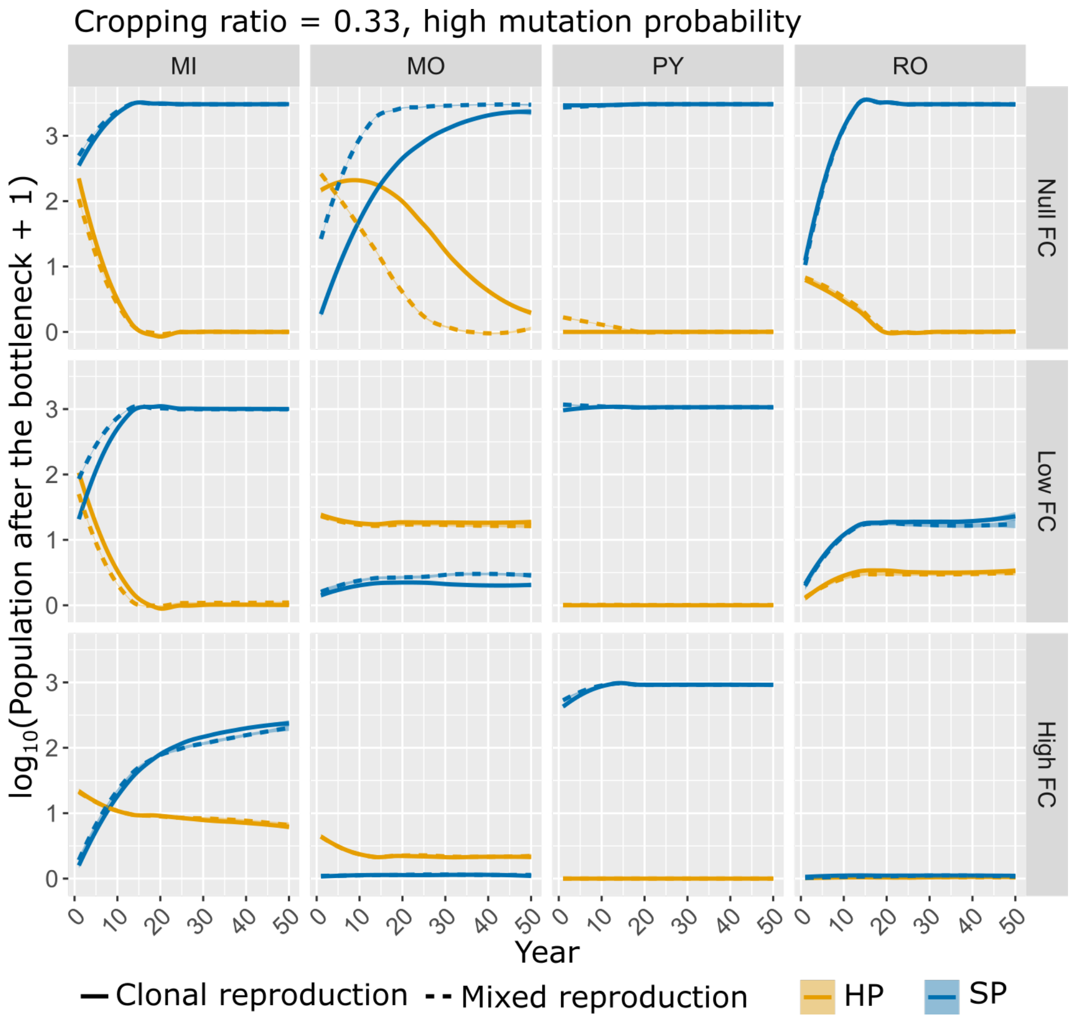


Figure S5: Population size of the superpathogen $SP_{tf}$ (in blue) and maximum number of heterogeneous parental pairs $HP_{tf}$ (in orange) in the landscape after the bottleneck. Curves represent populations dynamics across resistance deployment strategies (MIxture, MOsaic, ROtation and PYramiding) fitness costs and reproduction systems, at high mutation probability ($\tau = 10^{-4}$) and cropping ratio $\varphi_1 = 0.33$. Curves are based on the local polynomial regression fitting performed on simulations outputs. Shaded envelopes delimit the 2.5th and 97.5th percentiles.

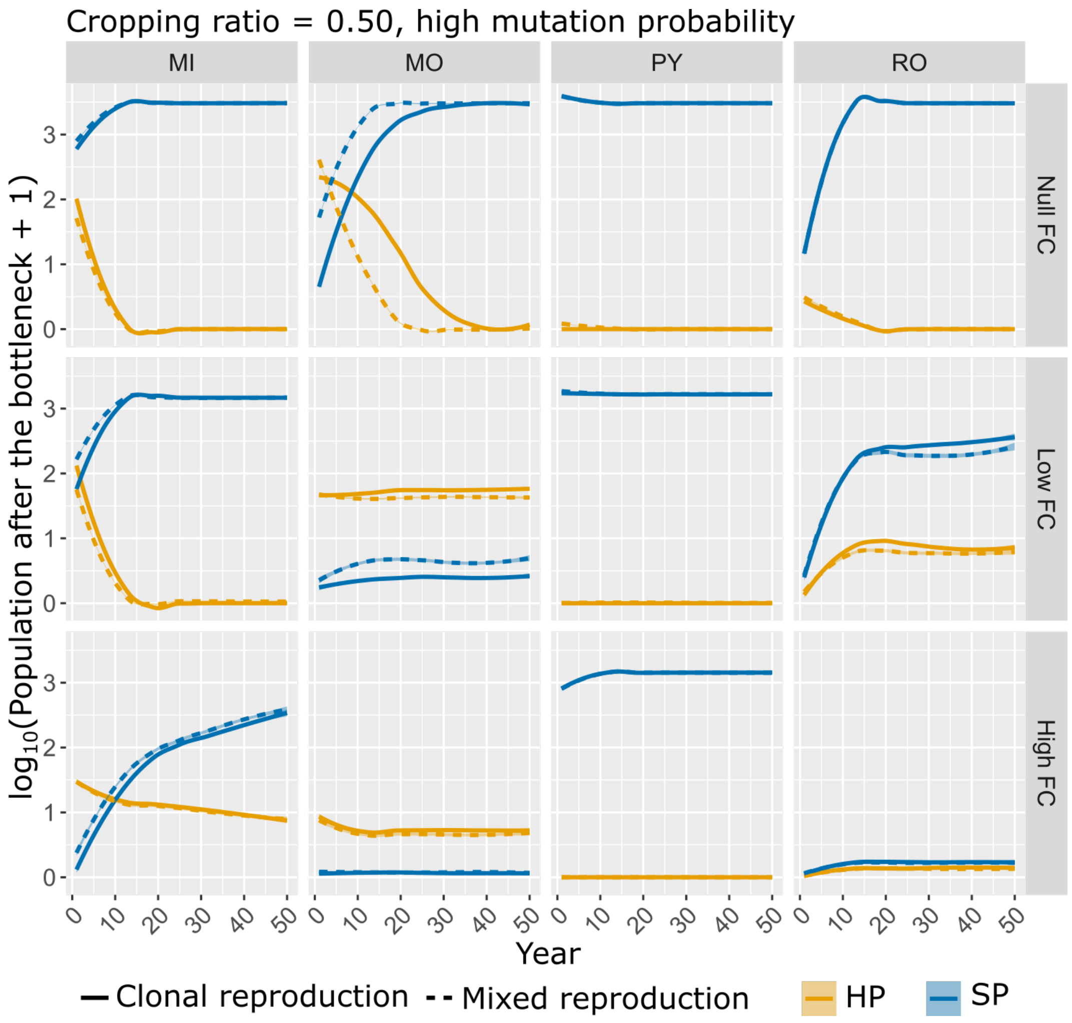


Figure S6: Population size of the superpathogen $SP_{t^f}$ (in blue) and maximum number of heterogeneous parental pairs $HP_{t^f}$ (in orange) in the landscape after the bottleneck. Curves represent populations dynamics across resistance deployment strategies (MIxture, MOsaic, ROtation and PYramiding) fitness costs and reproduction systems, at high mutation probability ($\tau = 10^{-4}$) and cropping ratio $\varphi_1 = 0.50$. Curves are based on the local polynomial regression fitting performed on simulations outputs. Shaded envelopes delimit the 2.5th and 97.5th percentiles.

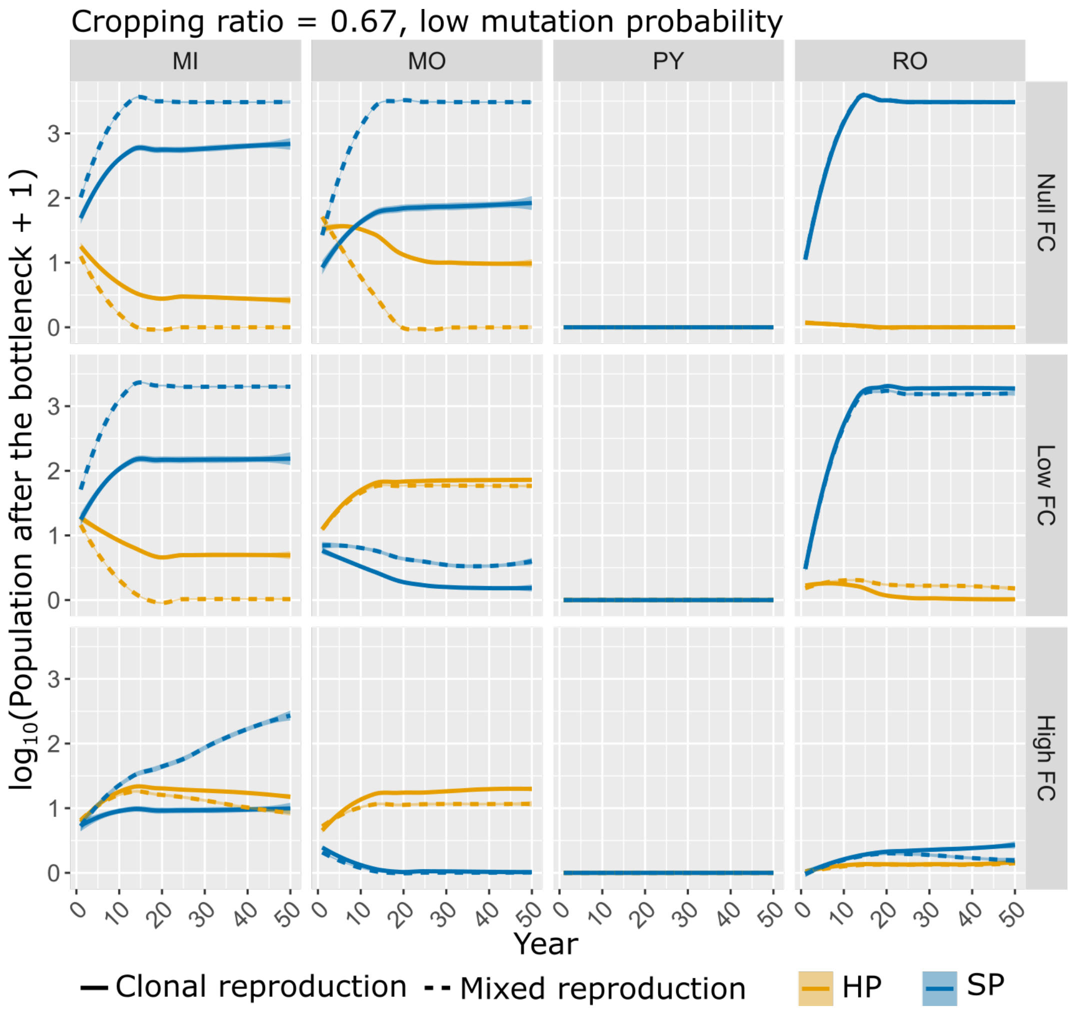


Figure S7: Population size of the superpathogen $SP_{tf}$ (in blue) and maximum number of heterogeneous parental pairs $HP_{tf}$ (in orange) in the landscape after the bottleneck. Curves represent populations dynamics across resistance deployment strategies (MIxture, MOsaic, ROtation and PYramiding) fitness costs and reproduction systems, at low mutation probability ($\tau = 10^{-7}$) and cropping ratio $\varphi_1 = 0.67$. Curves are based on the local polynomial regression fitting performed on simulations outputs. Shaded envelopes delimit the 2.5th and 97.5th percentiles.

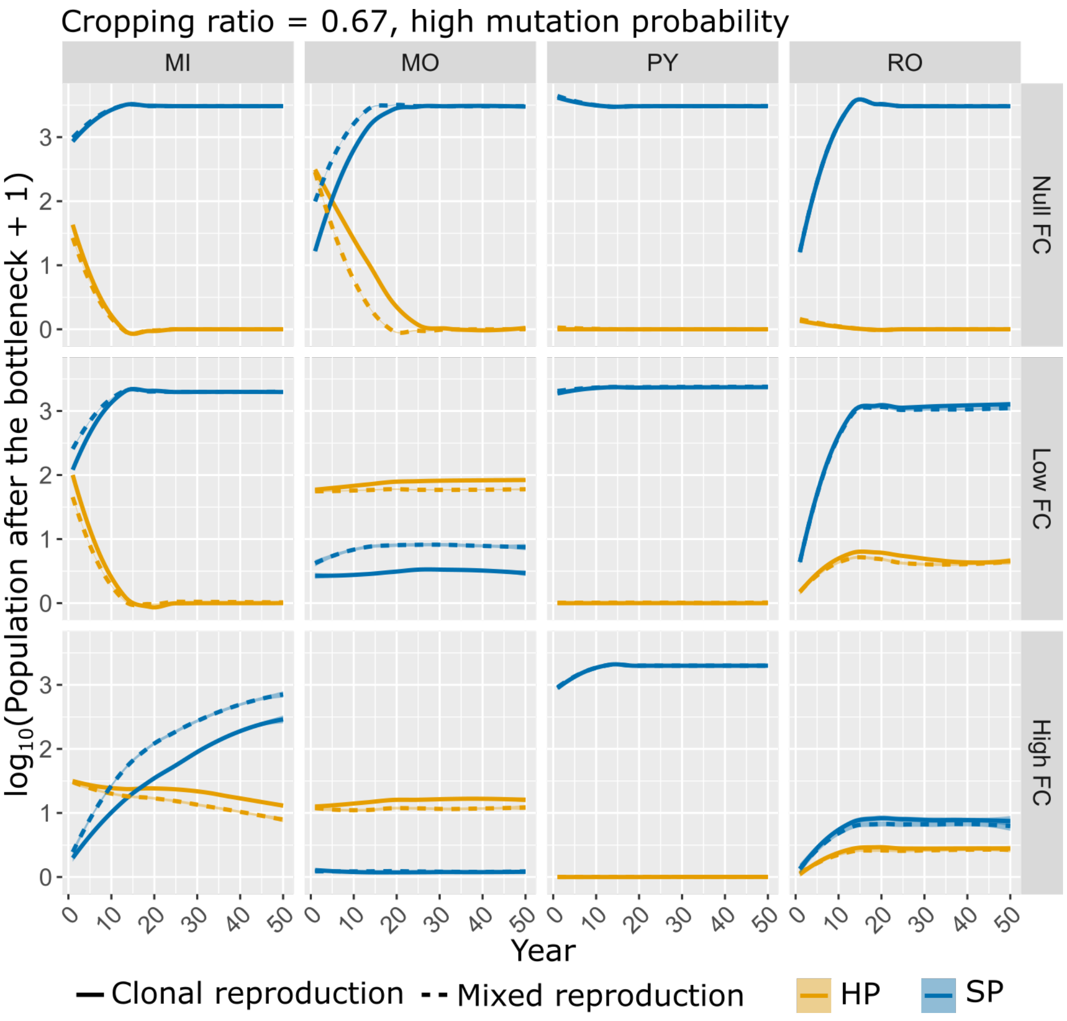


Figure S8: Population size of the superpathogen $SP_{tf}$ (in blue) and maximum number of heterogeneous parental pairs $HP_{tf}$ (in orange) in the landscape after the bottleneck. Curves represent populations dynamics across resistance deployment strategies (MIxture, MOsaic, ROtation and PYramiding) fitness costs and reproduction systems, at high mutation probability ($\tau = 10^{-4}$) and cropping ratio $\varphi_1 = 0.67$. Curves are based on the local polynomial regression fitting performed on simulations outputs. Shaded envelopes delimit the 2.5th and 97.5th percentiles.

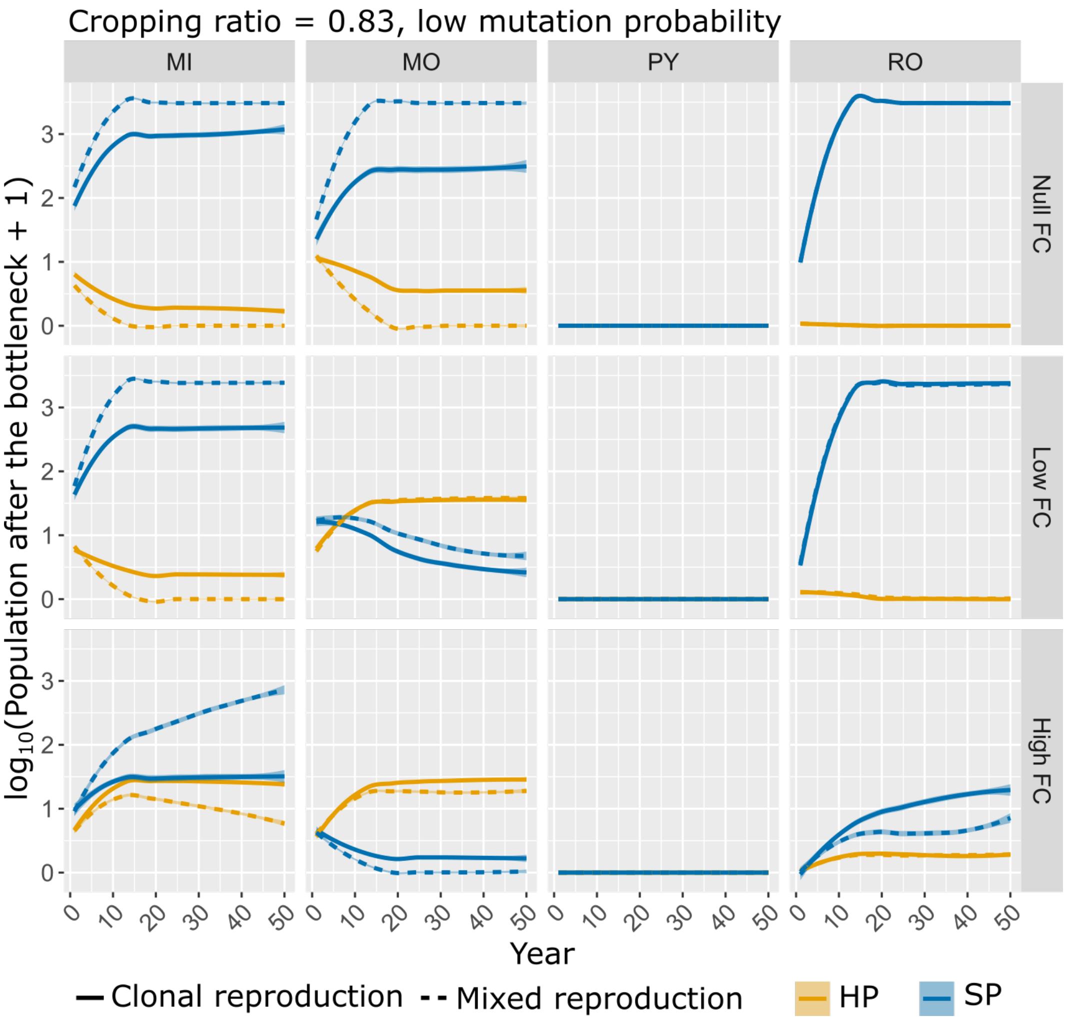


Figure S9: Population size of the superpathogen $SP_{tf}$ (in blue) and maximum number of heterogeneous parental pairs $HP_{tf}$ (in orange) in the landscape after the bottleneck. Curves represent populations dynamics across resistance deployment strategies (MIxture, MOsaic, ROtation and PYramiding) fitness costs and reproduction systems, at low mutation probability ($\tau = 10^{-7}$) and cropping ratio $\varphi_1 = 0.83$. Curves are based on the local polynomial regression fitting performed on simulations outputs. Shaded envelopes delimit the 2.5th and 97.5th percentiles.

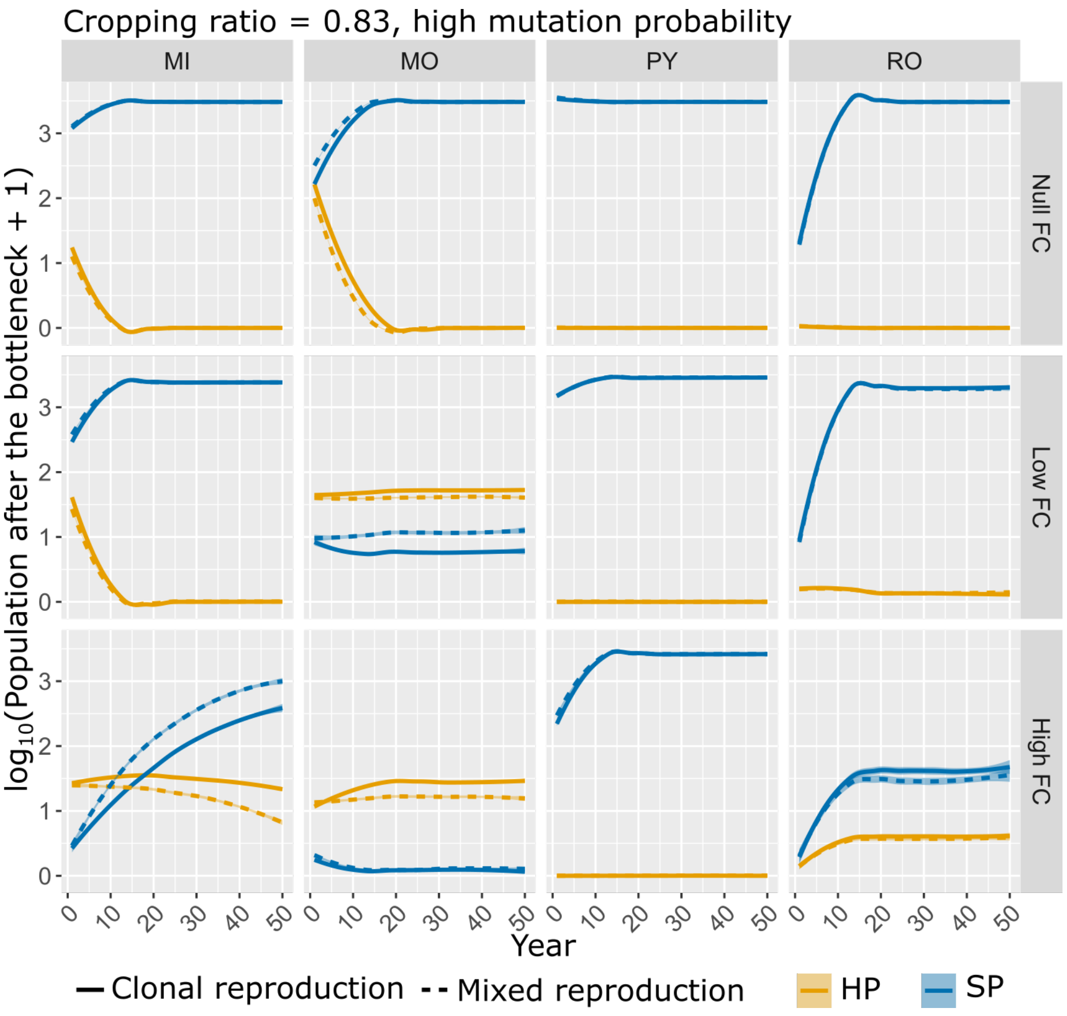


Figure S10: Population size of the superpathogen $SP_{tf}$ (in blue) and maximum number of heterogeneous parental pairs $HP_{tf}$ (in orange) in the landscape after the bottleneck. Curves represent populations dynamics across resistance deployment strategies (MIxture, MOsaic, ROtation and PYramiding) fitness costs and reproduction systems, at high mutation probability ($\tau = 10^{-4}$) and cropping ratio $\varphi_1 = 0.83$. Curves are based on the local polynomial regression fitting performed on simulations outputs. Shaded envelopes delimit the 2.5th and 97.5th percentiles.

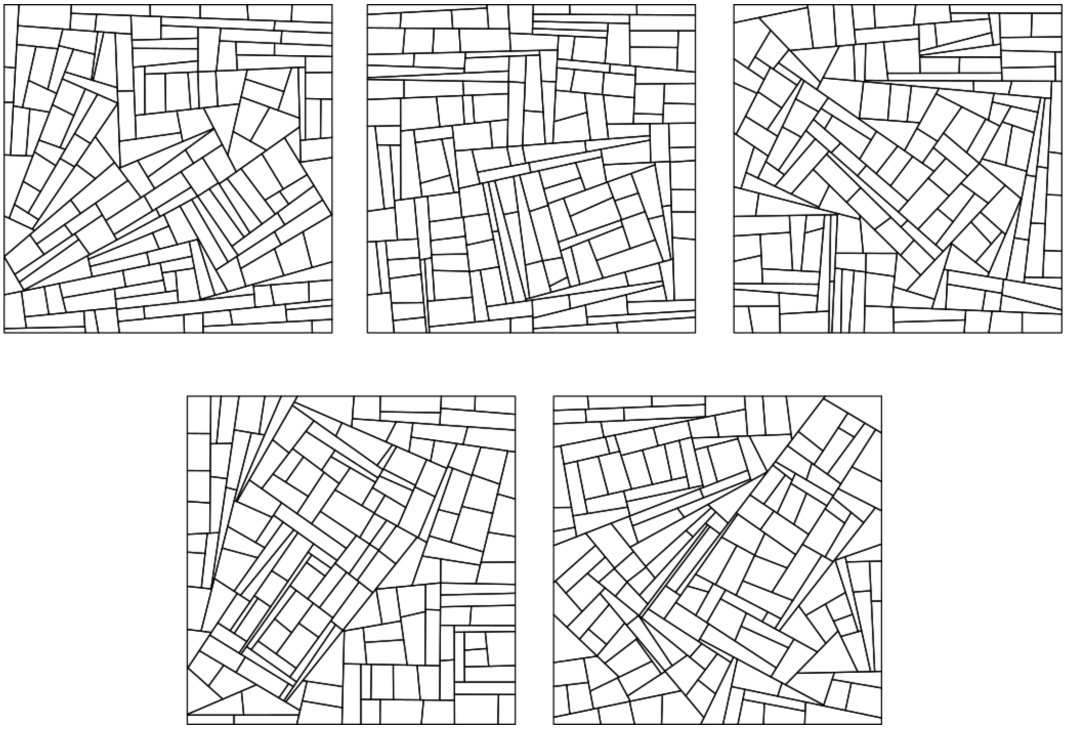

Figure S11: The five landscapes considered in the simulation plan. The landscapes account for 155, 154, 152, 153 and 156 fields, respectively. The total landscape area is 400 ha and the fields area ranges from 0.36 to 5.38 ha (mean: 2.60 ha).

### Note S1 Model equations

In the present work, the model is an adapted version of the one presented in Rimbaud et al. (2018), which simulates the clonal reproduction, spread and evolution of a pathogen in an agricultural landscape over multiple cropping seasons. Mainly, we introduce between-season sexual reproduction, characteristic of pathogens with a mixed reproduction system. We split the modelled cropping season into two distinct time periods: *i*) within cropping season, where multiple clonal reproduction events take place, and *ii*) between cropping seasons when a single sexual reproduction event may take place. The entire model is described in the following sections. Note that in the equations below only major resistance genes are considered. See Rimbaud et al. (2018) for details on model equations considering both major genes and quantitative resistance traits.

#### S1. 1 Host and pathogen demo-genetic dynamics within cropping season

The demo-genetic dynamics of the host-pathogen interaction are based on a HLIR structure ("healthy-latent-infectious-removed"). Thus, in the following, $H_{i,v,t}$, $L_{i,v,p,t}$, $I_{i,v,p,t}$, $R_{i,v,p,t}$, and $Pr_{i,p,t}$ respectively denote the number of healthy, latent, infectious and removed individuals (in this model, an "individual" is a given amount of plant tissue, and is referred to as a "host" hereafter for simplicity), and pathogen propagules in field $i$ ($i$=1,...,$J$), for cultivar $v$ ($v$=1,...,$V$), pathogen genotype $p$ ($p$=1,...,$P$) at time step $t$ ($t$=1,...,$T \times Y$). $T$ is the number of time steps in a cropping season and $Y$ the number of simulated years (*i.e.* cropping seasons). Since the host is cultivated, we assume there is no host reproduction, dispersal or natural mortality (leaf senescence near the end of the cropping season is considered as part of host harvest).

**Host growth.** Only healthy hosts (denoted as $H_{i,v,t}$) are assumed to contribute to growth of the crop. Thus, at each step $t$ during a cropping season, the plant cover of cultivar $v$ in field $i$ increases as a logistic function, and the new amount of healthy plant tissue is:

$$H_{i,v,t+1} = H_{i,v,t}\left[1 + \delta_v \times \left(1 - \frac{N_{i,v,t}}{K_{i,v}}\right)\right] \quad (1)$$

with $\delta_v$ the growth rate of cultivar $v$; $N_{i,v,t} = H_{i,v,t} + \sum_{p=1}^{P}\left(L_{i,v,p,t} + I_{i,v,p,t} + R_{i,v,p,t}\right)$ the total number of hosts in field $i$ for cultivar $v$ and at time $t$; and $K_{i,v} = A_i \times C_v^{max}$ the carrying capacity of cultivar $v$ in field $i$, which depends on $A_i$, the area of the field, and $C_v^{max}$, the maximal density for cultivar $v$. Note that when a mixture of several cultivars is present in the same field, decreased growth due to susceptible plants being diseased is not compensated by increased growth of resistant plants.

**Contamination of healthy hosts.** The healthy compartment ($H$) is composed of hosts which are free of pathogen propagules ($H^1$), as well as hosts contaminated (but not yet infected) by the arrival of such propagules ($H^2$). At the beginning of each step, all healthy hosts are considered free of propagules ($H^1$). Then at time $t$ in field $i$ and for cultivar $v$, the number of contaminable hosts (*i.e.* accessible to pathogen propagules, denoted as $H_{i,v,t}^{contaminable}$) depends on the proportion of healthy hosts ($H_{i,v,t}^1$) in the host population ($N_{i,v,t}$):

$$H_{i,v,t}^{contaminable} \sim Binomial\left(H_{i,v,t}^1; \pi\left(\frac{H_{i,v,t}^1}{N_{i,v,t}}\right)\right) \quad (2)$$

with $\pi(x) = \frac{1-e^{-\kappa x^{\sigma}}}{1-e^{-\kappa}}$, a sigmoid function with $\pi(0) = 0$ and $\pi(1) = 1$, giving the probability for a healthy host to be contaminated. Here, we assume that healthy hosts are not equally likely to be contacted by propagules, for instance because of plant architecture. Moreover, as the local severity of disease increases, eventually the probability for a single propagule to contaminate a healthy host declines due to the decreased availability of host tissue.

Following the arrival of propagules of pathogen genotype $p$ in field $i$ at time $t$ (denoted as $Pr^4_{i,p,t}$, see below), susceptible hosts become contaminated. The pathogen genotypes of these propagules are distributed among contaminable hosts according to their proportional representation in the total pool of propagules. Thus, for cultivar $v$, the vector describing the maximum number of contaminated hosts by each pathogen genotype (denoted as $[H^{maxConta}_{i,v,t}]_{p=1,...,P}$) is given by a multinomial draw:

$$[H^{maxConta}_{i,v,t}]_{p=1,...,P} \sim Multinomial\left(H^{contaminable}_{i,v,t};\left[\frac{Pr^4_{i,p,t}}{\sum_{p=1}^{P} Pr^4_{i,p,t}}\right]_{p=1,...,P}\right) \tag{3}$$

However, the number of deposited propagules ($Pr^4_{i,p,t}$) may be smaller than the maximal number of contaminated hosts ($H^{maxConta}_{i,v,p,t}$). Thus, the true number of hosts of cultivar $v$, contaminated by pathogen genotype $p$ in field $i$ at $t$ (denoted as $H^2_{i,v,p,t}$) is given by:

$$[H^1_{i,v,p,t} \rightarrow H^2_{i,v,p,t}] = min\left(H^{maxConta}_{i,v,p,t}; Pr^4_{i,p,t}\right) \tag{4}$$

**Infection.** Between $t$ and $t+1$, in field $i$, contaminated hosts ($H^2_{i,v,p,t}$) become infected (state L) with probability $e_{v,p}$, which depends on the maximum expected infection efficiency, $e_{max}$, and the interaction between host ($v$) and pathogen ($p$) genotypes:

$$[H^2_{i,v,p,t} \rightarrow L_{i,v,p,t+1}] \sim Binomial\left(H^2_{i,v,p,t}; e_{v,p}\right) \tag{5}$$

$$e_{v,p} = e_{max} \times \prod_{g=1}^{G} INF^g_{ig_g(p),mg(v)} \tag{6}$$

$INF^g$ represent the infectivity matrix for major gene $g$ which summarizes the possible interactions between potential host resistance genes ($mg$) and associated pathogen infectivity genes ($ig$), see Table 1 in the main text for an example.

**Latent period.** Infected hosts become infectious (state I) after a latent period (LI) drawn from a Gamma distribution (a flexible continuous distribution from which durations in the interval $[0; +\inf[$ can be drawn) parameterised with a minimal expected value, $\Gamma_{min}$, and variance, $\Gamma_{var}$:

$$(LI) \sim Gamma(\Gamma_{min}; \Gamma_{var}) \tag{7}$$

Note, the usual shape and scale parameters of a Gamma distribution, $\beta_1$ and $\beta_2$, can be calculated from the expectation and variance, exp and var, with: $\beta_1 = \frac{exp^2}{var}$ and $\beta_1 = \frac{var}{exp}$, respectively.

**Infectious period.** Finally, infectious hosts become epidemiologically inactive (*i.e.* they no longer produce propagules, thus are in state R, "removed") after an infectious period (IR) drawn from a Gamma distribution parameterised with maximal expected value, $\Upsilon_{max}$ and variance, $\Upsilon_{var}$, similar to the latent period:

$$(IR) \sim Gamma(\Upsilon_{max}; \Upsilon_{var}) \tag{8}$$

**Pathogen clonal reproduction.** In field $i$ at time $t$, infectious hosts associated with pathogen genotype $p$ produce a total number of propagules (denoted as $Pr^1_{i,p,t}$), drawn from a Poisson distribution with parameter $r_{max}$ corresponding to the maximal expected number of propagules produced by a single infectious host per time step:

$$Pr^1_{i,p,t} \sim Poisson(\sum_{v=1}^{V} r_{max}) \tag{9}$$

**Pathogen mutation.** The following algorithm is repeated independently for every potential infectivity gene g:

1. the pathotype (*i.e.* the level of adaptation with regard to major gene $g$, indexed by $q$; $q = 1, \ldots, Q_g$; with $Q_g = 2$ since the pathotype is either infective, or non-infective) of the pathogen propagules is retrieved from their genotype $p$;

2. propagules can mutate from pathotype $q$ to pathotype $q'$ with probability $m^g_{qq'}$ such as $m^g_{qq'} = \tau_g$ if $q' \neq q$ (hence $m^g_{qq} = 1 - \tau_g$ since $Q_g = 2$). Thus, in field $i$ at time $t$, the vector of the number of propagules of each pathotype arising from pathotype $q$ (denoted as $[M^g_{i,q,q',t}]_{q'=1,\ldots,Q_g}$) is given by a multinomial draw:

$$[M^g_{i,q,q',t}]_{q'=1,\ldots,Q_g} \sim Multinomial\left(Pr^1_{i,p,t}; [m^g_{qq'}]_{q'=1,\ldots,Q_g}\right) \tag{10}$$

3. the total number of propagules belonging to pathotype $q'$ and produced in field $i$ at time $t$ (denoted as $Pr^2_{i,q',t}$) is:

$$Pr^2_{i,q',t} = \sum_{q=1}^{Q_g} M^g_{i,q,q',t} \tag{11}$$

4. the new propagule genotype $p'$ is retrieved from its new pathotype ($q'$), and the number of propagules is incremented using a variable denoted as $Pr^3_{i,p',t}$.

In this model, it should be noted that the mutation probability $\tau_g$ is not the classic mutation rate (*i.e.* the number of genetic mutations per generation per base pair), but the probability for a propagule to change its infectivity on a resistant cultivar carrying major gene $g$. This probability depends on the classic mutation rate, the number and nature of the specific genetic mutations required to overcome major gene $g$, and the potential dependency between these mutations.

**Pathogen dispersal.** Propagules (both clonal and sexual, see below) can migrate from field $i$ (whose area is $A_i$) to field $i'$ (whose area is $A'_i$) with probability $\mu_{ii'}$, computed from:

$$\mu_{ii'} = \frac{\int_{A_i} \int_{A_{i'}} g(\|z' - z\|) dz dz'}{A_i} \tag{12}$$

with $\|z' - z\|$ the Euclidian distance between locations $z$ and $z'$ in fields $i$ and $i'$, respectively, and $g(\|z' - z\|) = \frac{(b-2)(b-1)}{2\pi a^2} \left(1 + \frac{\|z'-z\|}{a}\right)^{-b}$ the two-dimensional power law dispersal kernel of the propagules. The computation of $\mu_{ii'}$ probability is performed using the *CaliFloPP* algorithm (Bouvier et al., 2009). Thus, at time $t$, the vector of the number of propagules of genotype $p$ migrating from field $i$ to each field $i'$ (denoted as $[D_{i,i',p,t}]_{i'=1,\ldots,J}$) is:

$$[D_{i,i',p,t}]_{i'=1,\ldots,J} \sim Multinomial(Pr^3_{i,p,t}; [\mu_{i,i'}]_{i'=1,\ldots,J}) \tag{13}$$

and the total number of propagules arriving in field $i'$ at time $t$ (denoted as $Pr^4_{i',p,t}$) is:

$$Pr^4_{i',p,t} = \sum_{i=1}^{J} D_{i,i',p,t} \tag{14}$$

We consider that propagules landing outside the boundaries of the simulated landscape are lost (absorbing boundary condition), and there are no propagule sources external to the simulated landscape.

#### S1. 2 Host and pathogen demo-genetic dynamics between cropping seasons

**Seasonality.** Let $t^0(y)$ and $t^f(y)$ denote the first and last days of cropping season $y$ ($y = 1, \ldots, Y$), respectively. The plant cover in field $i$ for cultivar $v$ at the beginning of cropping season $y$ is set at $H_{i,v,t^0(y)} = A_i \times C^0_v \times \mathbb{I}_{v(i)=v}$, with $C^0_v$ the plantation density of cultivar $v$ and $\mathbb{I}_{v(i)}$ an indicative variable set at 1 when field $i$ is cultivated with cultivar $v$ and 0 otherwise. At the end of a cropping season, the host is harvested. We assume that the pathogen needs a green bridge to survive the off-season. This green bridge could, for example, be a wild reservoir or volunteer plants remaining in the field (*e.g.* owing to incomplete harvest or seedlings). The size of this reservoir imposes a bottleneck for the pathogen population. The number of remaining infected hosts in field

$i$ for cultivar $v$ and pathogen genotype $p$ (denoted by $I^*_{i,v,p,t^{f(y)}}$) at the end of the off-season is given by:

$$I^*_{i,v,p,t^{f(y)}} \sim Binomial(L_{i,v,p,t^{f(y)}}) + I_{i,v,p,t^{f(y)}}); \lambda) \tag{15}$$

with $\lambda$ the survival probability of infected hosts. Considering that those hosts produce propagules during their whole infectious period, we compute an equivalent number of infectious hosts by multiplying $I^{eq}_{i,v,p,t^{f(y)}} = \Upsilon_{max} \times I^*_{i,v,p,t^{f(y)}}$. The remaining hosts $I^{eq}_{i,v,p,t^{f(y)}}$ produce clonal and sexual propagules. Clonal propagules can mutate exactly as happens during the cropping season. The production of propagules through sexual reproduction and the possible genetic recombination are detailed in the following section. Propagules, either clonal or sexual, produced between cropping seasons are uniformly released throughout the following cropping season, constituting the primary inoculum.

**Pathogen sexual reproduction.** In field $i$, the pool of infectious hosts associated to the same cultivar $v$, $I_{i,v,p,t^{f(y)}}$, undertakes sexual reproduction. Two parental infectious hosts, respectively infected by pathogens $Par_1$ and $Par_2$ are randomly sampled without replacement from the pool of infectious hosts. The couple $c = \{Par_1; Par_2\}$ produce $P^{sex}_{v,c}$ propagules, drawn from a Poisson distribution whose expectation is the sum of the number $r_{max}$ of propagules produced by each of the parental infectious hosts:

$$P^{sex}_{v,c} \sim Poisson(r_{expv,c} = r_{max} + r_{max} = 2 \times r_{max}) \tag{16}$$

Then the genotype of each propagule is retrieved from parental genotypes: the genotype at every locus $g$ is randomly sampled between one of the two parents $\{Par_1; Par_2\}$. For example, assuming that parental infection $Par_1$ carries infectivity genes to resistance gene $R_1$ (which corresponds to a genotype "10") and parental infection $Par_2$ carries the infectivity genes to resistance $R_2$ (genotype "01"), the resulting propagule genotype could be either the same as one of the two parents, or a superpathogen genotype "11", or a wild-type genotype "00". This process is iterated for all the pairs $c$ = 1,..., $C$ of infectious hosts associated to all the cultivars $v$ = 1,..., $V$ in a given field $i$, resulting in a total number of sexual propagules:

$$P^{sex}_i = \sum_{v=1}^{V} \sum_{c=1}^{C} P^{sex}_{v,c} \tag{17}$$

#### S1. 3 Initial conditions

At the beginning of a simulation, healthy hosts are planted in each field. The initial pathogen population is assumed to be totally non-adapted to the resistance genes, and is only present in susceptible fields with probability $\Phi$. Then the initial number of infectious hosts in these fields is:

$$I_{i,v=1,p=1,t=1} \sim Binomial(H_{i,v=1,t=1}; \Phi) \tag{18}$$

### Note S2 Model parameterisation to downy mildew

This section details the parameterisation of the model to approximate the causal agent of downy mildew disease of grapewine, caused by the oomycete *Plasmopara viticola.*

#### S2. 1 Pathogen dispersal

Based on the results of previous empirical and modelling studies (Ojiambo et al., 2017; Mundt et al., 2009), the power-law has a good ability to predict the dispersal of aerially transmitted pathogens, assuming it is isotropic (*i.e.* uniform in all directions). We consequently use this function in our model:

$$g(\|z' - z\|) = \frac{(b-2)(b-1)}{2 \cdot \pi \cdot a^2} \left(1 + \frac{\|z' - z\|}{a}\right)^{-b} \tag{19}$$

where $a > 0$ is a scale parameter and $b > 2$ determines the weight of the dispersal tail. The expected dispersal distance is given by: $\mu_{exp} = \frac{2a}{(b-3)}$.

To our knowledge, there are no studies so far estimating the dispersal kernel, or at least an average dispersal distance, for *P. viticola* at a regional scale. For this reason, we decided to keep the same values reported in Rimbaud et al. (2018) for parameters of the dispersal kernel. Therefore, we calibrated $a$ and $b$ of our power-law function in such a way that the mean dispersal distance of a *P. viticola* spore is $\mu_{exp}$= 20 m, which represents 1% of landscape length. Among the possible solutions we arbitrarily chose $a = 40$ and $b = 7$. As soon as any information about the dispersal ability of *P. viticola* becomes available, these parameters will be easily updated accordingly.

#### S2. 2 Infection rate

In an experiment on the influence of environmental conditions on sporulation of *Plasmopara viticola* lesions, the probability that the inoculum drops deposited on a leaf triggers an infection was estimated to lie between 0.10 and 0.90, depending on the temperature and humidity (Caffi et al., 2016). In the present study, given that the propensity of a leaf to be infected is explicitly modelled by a sigmoid contamination function, the maximal expected infection rate was set at $e_{max} = 0.90$.

#### S2. 3 Sporulation rate

The real number of spores produced daily by a lesion and effectively dispersed to a leaf where they may trigger an infection, often summarised by 'effective sporulation rate' and denoted as $r_{max}$ here, is extremely complicated to estimate. As an indication, the basic reproductive number for a pathogen (*i.e.* the theoretical number of secondary infections from a single infectious host (Anderson and May, 1992)) would be $R_{0max} = e_{max} \times \Upsilon_{max} \times \mathrm{r}_{max}$. The $R_0$ of *P.*

*viticola* have been estimated to vary between 0.1-50 (Rossi et al., 2009). Thus, in order to simulate a reasonably aggressive pathogen, and since very few data were available to quantify $r_{max}$ compared to other parameters, $r_{max}$ has been adjusted to be 2 spores.day$^{-1}$, thus $R_{0max} = 25.05$ (mean value of the range proposed by Rossi et al., 2009).

#### S2. 4 Duration of the latent and sporulation periods

We found 13 estimates from 12 studies of *P. viticola* which included information on the duration of the latent period; similarly, there were 6 estimates from 5 studies with data on the sporulation period. When available, standardised conditions in growth chambers were preferred over field data to facilitate comparison between studies. A Gamma distribution was fit to the data using a maximum likelihood approach (figures S12 and S13). From this analysis, the parameters associated with latent and sporulation periods were estimated as: $\gamma_{min} = 7$; $\gamma_{var} = 8$; $\Upsilon_{max} = 14$ and $\Upsilon_{var} = 22$.

Table S1: Available data on the duration of latent and sporulation periods for downy mildew caused by *P. viticola* (and *formae speciales*). In the following studies, latent and sporulation periods were determined on fully susceptible hosts under controlled condition (cyan cells), in field studies (green cells) or they are derived by biological evidences or other modelling frameworks (yellow cells).

| Reference | Organism | Disease | Latent Period (days) | | | Sporulation perido (days) | | |
|---|---|---|---|---|---|---|---|---|
| | | | min | max | mean | min | max | mean |
| Delmas et al. (2016) | *P. viticola* | Downey mildew | 2.8 | 3.5 | 3.15 | | | |
| Lalancette et al. (1988) | *P. viticola* | Downey mildew | 7 | 12 | 10 | | | |
| Bove et al. (2020) | *P. viticola* | Downey mildew | 6 | 10 | 8 | 15 | 20 | 17.5 |
| Caffi et al. (2013) | *P. viticola* | Downey mildew | 4 | 23 | 11.3 | 2 | 10 | 6.7 |
| Lalancette (1988) | *P. viticola* | Downey mildew | | | 7 | | | |
| Kennelly et al. (2005) | *P. viticola* | Downey mildew | 7 | 10 | 8.5 | | | |
| Mouafo-Tchinda et al. (2020) | *P. v. riparia* | Downey mildew | 5.75 | 6.25 | 6 | | | |
| Mouafo-Tchinda et al. (2020) | *P. v. aestivalis* | Downey mildew | 3.75 | 4.25 | 4 | | | |
| Rumbolz et al. (2002) | *P. viticola* | Downey mildew | 5 | 6 | 5.5 | | | |
| Boso and Kassemeyer (2008) | *P. viticola* | Downey mildew | | | 5 | | | |
| Rossi et al. (2009) | *P. viticola* | Downey mildew | | | 8 | | | 15 |
| Clark and Spencer-Phillips (2004) | *P. viticola* | Downey mildew | 3 | 24 | 13.5 | | | |
| Angelotti et al. (2017) | *P. viticola* | Downey mildew | 5 | 9 | 7 | | | |
| Bove et al. (2019) | *P. viticola* | Downey mildew | | | | 6 | 30 | 18 |
| Kennelly et al. (2007) | *P. viticola* | Downey mildew | | | | 6 | 18 | 12 |

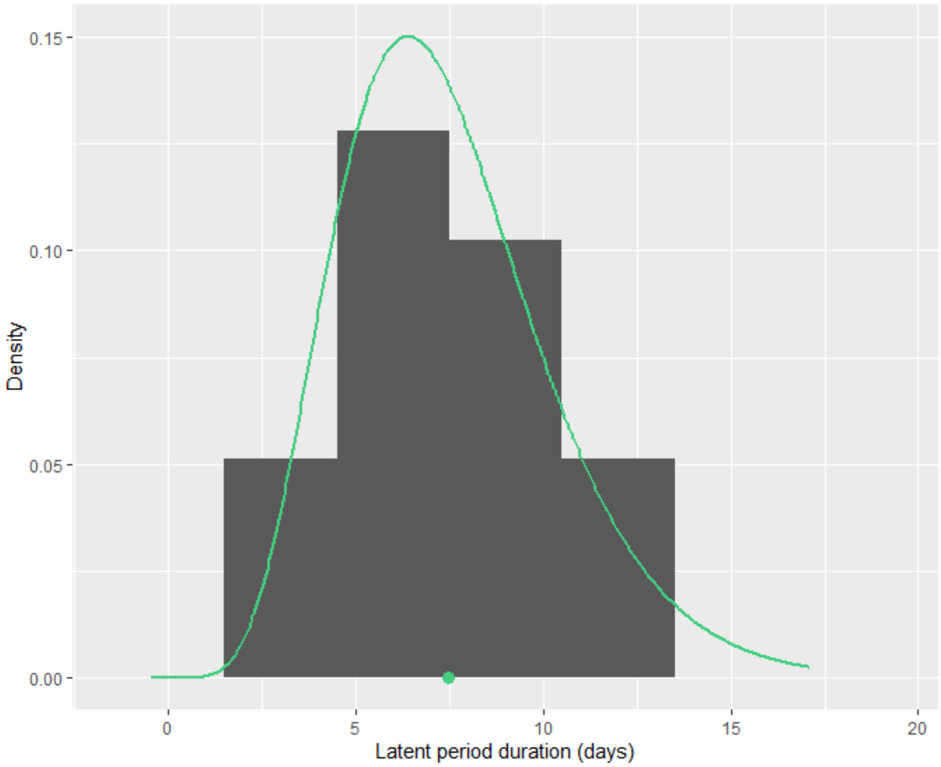


Figure S12: Distribution of the latent period duration of downy mildew caused by *Plasmopora viticola*. Raw data were obtained from previous studies (see Table S1). Green curve is Gamma distribution estimated from the data through maximum likelihood (function fitdistr of the R package MASS, v 7.3-53.1); dot indicate the mean latent period (7.46 days). The variance is 8.02 days.

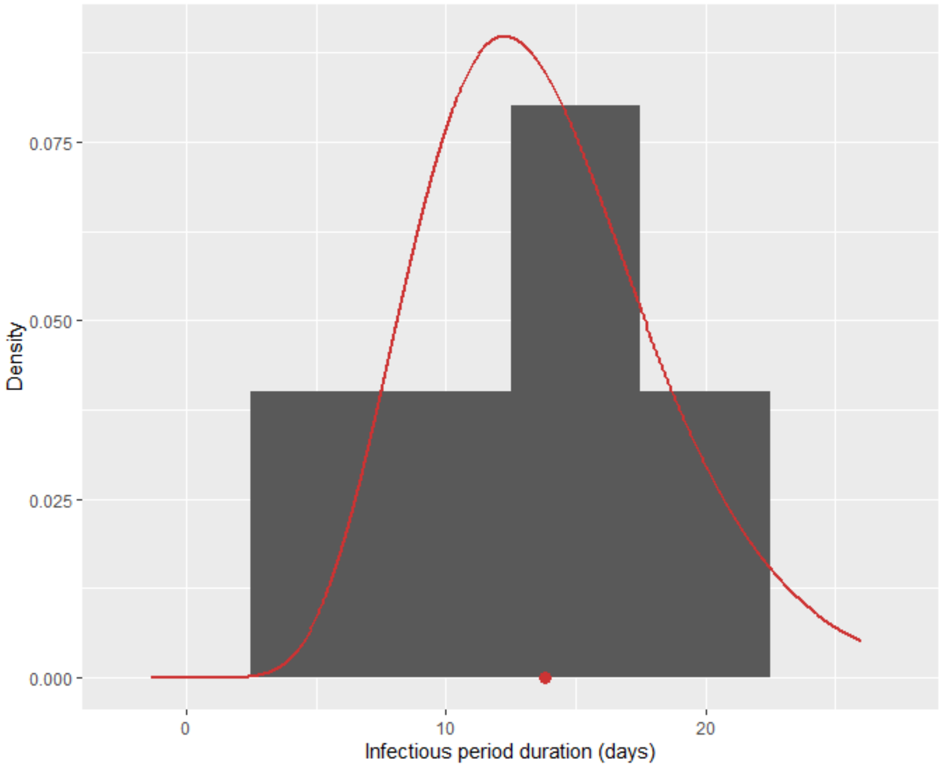


Figure S13: Distribution of the infectious period duration of downy mildew caused by *Plasmopora viticola*. Raw data were obtained from previous studies (see Table S1). Red curve is Gamma distribution estimated from the data through maximum likelihood (function fitdistr of the R package MASS, v 7.3-53.1); dot indicate the mean infectious period (13.84 days). The variance is 21.82 days.

#### S2. 5 Contamination function

We assume that susceptible plants become less easily contaminated when the local disease level increases by using the following sigmoid function for $\pi(x)$:

$$\pi(x) = \frac{1 - exp(-\kappa x^{\sigma})}{1 - exp(-\kappa)} \tag{20}$$

The parameters of the sigmoid curve ($\kappa = 5.33, \sigma = 3$) have been parameterised as in previous works (Papaïx et al., 2014b,a, 2015; Rimbaud et al., 2018), such as $\pi(0) = 0, \pi(1) = 1$, and the inflexion point is located at $x_0 = [(\sigma - 1)/(\kappa\sigma)]^{1/\sigma} \approx 0.5$. The use of a sigmoid function instead of a linear function implies that the contamination of a susceptible plant is easier when the proportion of susceptible plants is higher than 50%, and harder when the proportion of susceptible plants is lower than 50%.

### Note S3 Calculation of the threshold for pathogen establishment considering sexual reproduction

In this work, we defined a threshold (denoted by N hereafter) for the number of infections of resistant hosts (considering a constant and infinite host population size) by mutant pathogens with genotype $m_1$, above which they are unlikely to go extinct. The time passed between the beginning of a simulation and the moment the number of infections of resistant hosts by mutant pathogens $m_1$ exceeds the threshold N is considered the time of genotype $m_1$ establishment.

The spread of the mutant genotype $m_1$ across years from two parental infections of a resistant host (for subsequent sexual reproduction) requires surviving the bottleneck imposed by host harvest (denoted by event $Surv_I$), the production of at least one propagule with genotype $m_1$ via sexual reproduction (denoted by the event $Rec_I|Surv_I$) and the infection of new resistant hosts in the next cropping season (denoted by event $Inf_I|Surv_I, Rec_I$). Thus the probability of spread of the mutant genotype $m_1$ across years (event $Inf_I$) is:

$$P(Inf_I) = P(Surv_I) \times P(Rec_I|Surv_I) \times P(Inf_I|Surv_I, Rec_I) \tag{21}$$

The probability for a single infection to survive the bottleneck imposed by host harvest and the off-season corresponds to the off-season survival probability $\lambda$. However, as sexual reproduction takes place, we need, at least, that two infections survive the bottleneck. In a population of N infections, the probability of *i*) none of the infections survives the bottleneck OR *ii*) just one infection survives the bottleneck is :

$$P(Ext) = (1-\lambda)^N + \lambda(1-\lambda)^{N-1} \tag{22}$$

Consequently, the probability that at least two infections survives the bottleneck is:

$$P(Surv_I) = 1 - [(1-\lambda)^N + \lambda(1-\lambda)^{N-1}] \tag{23}$$

We now assume that just two infections survive the bottleneck and sexually reproduce. The probability for a single propagule produced by the couple to have the same genotype $m_1$ of the parental infection $p_1$ is equal to 1 if the other parental infection $p_2$ has genotype $m_1$, too. By contrast if the parental infection $p_2$ has a genotype different from $p_1$, the probability for a single propagule to have genotype $m_1$ is:

$$P(Rec|m_1) = \prod_{g=1}^{G} \frac{1}{ig_g} \tag{24}$$

Where $G$ is the number of resistance genes considered and $ig_g$ is the number of level of aggressiveness for the resistance gene g (=2 for major genes, higher than 2 for QTLs). Note that in equation (4) we are considering the worst case scenario, that is the two parental infections have different aggressiveness for each resistance gene. If the parental infections $p_1$ and $p_2$ have the same level of aggressiveness for some resistance genes, $P(Rec|m_1)$ would be higher than that in equation (4). In our case, we consider two (G=2) major resistance genes (ig=2), then $P(Rec|m_1) = 0.25$. Consequently, the probability that the produced propagule does not inherit the genotypes $m_1$ from parent $p_1$ is $1 - \prod_{g=1}^{G} \frac{1}{ig_g} = 0.75$. Since two parental infections produces a total of $2 \times r_{max} \times \Gamma_{max}$ propagules, the probability that none of the propagules inherit the genotype $m_1$ is $P(Ext_{REC}) = 0.75^{(2 \times r_{max} \times \Gamma_{max})}$. Given the values $r_{max} = 2$, $\Gamma_{max} = 14$ (parameters for downy mildew), we have $P(Ext_{REC}) \approx 1 \times 10^{-7}$. Consequently, the probability of production of at least one propagule with genotype $m_1$ via sexual reproduction is:

$$P(Rec_I|Surv_I) = 1 - P(Ext_{REC}) \approx 1 \tag{25}$$

Concerning the probability of infecting a new resistant hosts the next cropping season ($P(Inf_I|Surv_I, Rec_I)$), the probability for a single propagule with genotype $m_1$ (completely adapted to the resistant host population) to $i$) stay mutant, that is, it does not incur reverse mutations (from mutant $m_1$ to wild type) and $ii$) infect a resistant host is given, respectively, by $(1 - \tau_g)$, where $\tau_g$ is the mutation probability for the infectivity gene $g$, and $e_{max}$, which is the infection rate. Then, the probability for the propagule to mutate and/or not infect any host is $1 - (1 - \tau_g)e_{max}$. Note that we assumed an infinite host population, which allows to neglect the probability to disperse outside the field and the competition with other pathogen genotypes. Two parental infections produce via sexual reproduction a total of $2 \times r_{max} \times \Gamma_{max}$ spores, which have $m_1$ genotype with probability $P(Rec_I|m_1) = 0.25$. Consequently, the probability that none of these propagules stays mutant and infects a host in the next cropping season is $P(Ext_{Pr}) = [1 - (1 - \tau_g)e_{max}]^{2 \times r_{max} \times \Gamma_{max} \times P(Rec_I)}$. Given the value $\tau_g = 10^{-4}$, $e_{max} = 0.9$, $r_{max} = 2$ and $\Gamma_{max} = 14$, $P(Ext_{Pr}) \approx 1 \times 10^{-14}$. Consequently, the probability of infection of new resistant hosts by propagules with genotype $m_1$ is:

$$P(Inf_I|Surv_I, Rec_I) = 1 - P(Ext_{Pr}) \approx 1 \tag{26}$$

The combination of equations (1), (3), (5) and (6) gives:

$$P(Inf_I) = P(Surv_I) \tag{27}$$

Then, the probability of extinction of N infections is as in equation (2).

Above N=50,000 infections, the probability of extinction is less than 1%. Therefore, we chose this threshold to define the time to mutant pathogen establishment.
